# A regulatory circuit involving the NADH dehydrogenase-like complex balances C_4_ photosynthetic carbon flow and cellular redox in maize

**DOI:** 10.1101/2023.02.23.529632

**Authors:** Qiqi Zhang, Shilong Tian, Genyun Chen, Qiming Tang, Yijing Zhang, Andrew J. Fleming, Xin-Guang Zhu, Peng Wang

## Abstract

- C_4_ plants typically operate a CO_2_ concentration mechanism from mesophyll (M) cells into bundle sheath (BS) cells. NADH dehydrogenase-like (NDH) complex is enriched in the BS cells of many NADP-ME type C_4_ plants, and is more abundant in C_4_ than in C_3_ plants, but to what extent it is involved in the CO_2_ concentration mechanism remains to be experimentally investigated.
- We created maize and rice mutants deficient in NDH function, and used a combination of transcriptomic, proteomic, and metabolomic approaches for comparative analysis.
- Considerable decrease in growth, photosynthetic activities, and levels of key photosynthetic proteins were observed in maize but not rice mutants. However, gene expression for many cyclic electron transport and Calvin-Benson cycle components plus BS specific C_4_ enzymes, was up-regulated in maize mutants. Metabolite analysis of the maize *ndh* mutants revealed increased NADPH/NADP ratio, as well as malate, RuBP, FBP, and photorespiration components.
- We suggest that by optimizing NADPH and malate levels, adjusting NADP-ME activity, NDH functions to balance metabolic and redox states in the BS cells of maize, coordinating photosynthetic gene expression and protein content, thus directly regulating the carbon flow in the two-celled C_4_ system of maize.

## Introduction

Plants use light energy to eventually drive CO_2_ assimilation into carbohydrates through photosynthesis. The organization and operation of the photosynthetic apparatus are finely regulated at different levels, and a deeper understanding of them would benefit crop improvement. Rice and wheat are the most widely consumed food crops, but the C_3_ photosynthesis in these plants does not operate in an efficient way, with the assimilation of CO_2_ considerably compromised due to photorespiration (Bauwe *et al*., 2010; Raines, 2011). Crops such as maize and sorghum have evolved a distinct and more efficient process, C_4_ photosynthesis. Typical C_4_ leaves display a special Kranz anatomy: vascular bundles of C_4_ maize leaves are sequentially surrounded by functionally differentiated bundle sheath (BS) cells and mesophyll (M) cells, which contain chloroplasts with different ultrastructure and a specific distribution of metabolic enzymes. In C_4_ plants, CO_2_ is concentrated through intercellular transport of C_4_ acids (such as malate and aspartate) from M cells where CO_2_ is captured via PEP carboxylase, to the BS cells where Rubisco is present. The C_4_ acids are decarboxylated in the BS cells, raising CO_2_ concentration around Rubisco, decreasing photorespiration, and thus achieving more efficient assimilation of CO_2_ and improved photosynthesis (Furbank, 2017; von Caemmerer *et al*., 2017).

One distinctive feature of C_4_ BS chloroplasts maybe linked to a strong ability to perform cyclic electron transport (CET). CET pathways are mainly mediated by the chloroplast NADH dehydrogenase-like complex (NDH), and by the proton gradient regulatory proteins PGR5 and PGRL1 (Johnson, 2011; Shikanai, 2014; Suorsa, 2015; Yamori *et al*., 2016). The NDH complex binds to PSI and is specifically distributed in the non-stacking regions of thylakoids (Peng *et al*., 2008, 2009, 2011; Kouřil *et al*., 2014; Yadav *et al*., 2017). By mediating CET from the receptor side of photosystem I (PSI) to plastoquinone (PQ), an NDH-dependent proton gradient is established across the thylakoid membrane to drive ATP synthesis (Wang *et al*., 2006; Ishikawa *et al*., 2016b; Strand *et al*., 2017). In C_3_ plants, CET is not prominent under normal conditions, and CO_2_ assimilation is mainly driven by linear photosynthetic electron transport (LET). However, under environmental stresses, such as high temperature, high light and drought, LET is suppressed, while CET via NDH is often up-regulated. This helps to decrease oxidative damage and to replenish ATP (Burrows *et al*., 1998; Endo *et al*., 1999; Horváth *et al*., 2000; Li *et al*., 2004; Wang *et al*., 2006; Ishikawa *et al*., 2008a; Yamori *et al*., 2011; Suorsa *et al*., 2015). In the C_4_ plant maize, BS chloroplasts lack the stacking grana thylakoids rich in photosystem II (PSII), and instead possess a large number of non-stacking stromal thylakoids rich in PSI and NDH complex (Majeran and van Wijk, 2009; Ermakova *et al*., 2020). The C_4_ plant and cell-specific up-regulation of NDH was confirmed at both transcript and protein levels (Kubicki *et al*., 1996; Darie *et al*., 2005, 2006; Takabayashi *et al*., 2005; Majeran *et al*., 2008; Friso *et al*., 2010; Li *et al*., 2010; Bräutigam *et al*., 2011). The functional significance of the enriched NDH content in C_4_ BS cells has aroused great interest.

It has been noted that NDH content is higher in a variety of C_4_ leaves (both NADP-ME and NAD-ME types) than in C_3_ leaves, leading to the idea that it offers increased ATP generation required by the C_4_ pathway (Ishikawa *et al*., 2016b). As evidence, studies on several C_4_ plants have shown that the NDHH subunit is increased in cell types requiring additional ATP for CO_2_ concentration (e.g., BS cells of NADP-ME type and M cells of NAD-ME type leaves) (Takabayashi *et al*., 2005). In some C_4_ species only NDH-mediated CET is enhanced, while in others both NDH and PGR5-PGRL1 pathways are enhanced (Munekage *et al*., 2010; Nakamura *et al*., 2013). As highlighted by the involvement of NDH but not PGR5 in the CO_2_ concentration mechanism (CCM) of cyanobacteria (Price *et al*., 2013; Long *et al*., 2016), whether NDH plays a more specific role apart from ATP production in the C_4_ pathway is still unknown. Further clue may be related with the involvement of NDH-CET in NADPH turnover and redox adjustment. Since various photosynthetic enzymes that take part in light reactions and carbon reactions are regulated via redox components, understanding the different redox needs in the M and BS chloroplasts of C_4_ plants is crucial for engineering efficient C_4_ photosynthesis (Turkan et al., 2018). Particularly, NADP-malic enzyme (NADP-ME) catalyzes the oxidative decarboxylation of malate to yield CO_2_ and pyruvate with the concomitant production of NADPH (Drincovich et al. 2001), being the main source of reducing power for BS chloroplasts in maize. For C_4_ photosynthesis to efficiently concentrate CO_2_ in BS chloroplasts, the NADP-ME decarboxylation and Rubisco carboxylation rates must be coordinated. Besides having acquired an enriched BS expression, the C_4_-type NADP-ME displays a particular redox modulation in maize (Alvarez et al., 2012). Whether or how NDH-CET, as well as the ferredoxin/thioredoxin system, are related to this modulation greatly attract our interest.

Several groups have taken a molecular genetic approach to understand the relationship between NDH-mediated CET and C_4_ photosynthesis. Studies have found that mutations in the NDHN or NDHO subunits of maize lead to increased photorespiration and reduced carbon assimilation under high-light and saturated CO_2_ conditions (Peterson *et al*., 2016). In the C_4_ species *Flaveria bidentis*, knockout of the NDHN or NDHO subunit, but not PGR5 nor PGRL1, resulted in slow plant growth and decreased CO_2_ assimilation (Ishikawa *et al*., 2016a; Ogawa *et al*., 2023). These studies indicate that the function of NDH is crucial for photosynthesis and growth of C_4_ plants, but the mechanism underlying the phenotype remains to be further studied. In this study, we took a reverse genetic approach and created mutants of selected NDH subunits (NDF6 and NDHU subunits), both in a C_4_ crop (maize) and a C_3_ crop (rice), then used a combination of transcriptomic, proteomic, and metabolomic approaches to identify the mechanism by which loss of NDH activity in C_4_ leaves leads to decreased CO_2_ assimilation and growth. Our data highlight the importance of BS-localized NDH activity in maize for the balancing of carbon flux and redox state between the BS and MS cells, in addition to the generally accepted role of providing ATP. The necessity of this regulation is most prominent in the BS cells, which undergo continuous import of malate and generation of NADPH as an essential part of the C_4_ mechanism.

## Materials and Methods

### Plant growth conditions and trait measurement

Maize plants were grown in 6 L pots with a mix of 60% peat soil and 20% vermiculite. Rice plants were grown in 6 L pots with field soil. The plants were grown in environmentally controlled phytotron room [27 °C in day and 25 °C at night, 600 µmol photons m^−2^ s^−1^ photosynthetic photon flux density (PPFD), 16 h light and 8 h dark photoperiod, and ∼70% relative humidity] with normal fertilizer application.

To estimate leaf total chlorophyll content, a SPAD 502 Plus Chlorophyll Meter (Spectrum Technologies) was used. Plant height was measured from the soil surface to the collar of the youngest fully expanded leaf.

### Monitoring chlorophyll fluorescence and P700 redox

Post-illumination increase of chlorophyll fluorescence were monitored after turning off the actinic light (200 µmol photons m^−2^ s^−1^, white light, for 180 sec), using a PAM chlorophyll fluorometer (PAM101, Walz, Germany) attached with emitter-detector assembly and 101ED unit, as previously described (Wang *et al*., 2006). Fv/Fm, ETR, and NPQ were measured using a PAM-2000 chlorophyll fluorometer (Walz, Germany) according to standard procedures.

The redox kinetics of P700 was recorded using a Dual-PAM-100 instrument (Walz, Germany). Before measurements, the leaves were dark-acclimated for 20 min. Far-red light (30 sec) was applied to start the measurement, and the initial rate of P700^+^ reduction following termination of far-red light was calculated.

### Photosynthetic gas exchange measurements

The photosynthetic rate of phytotron-grown plants 60 days after transplanting was measured using LI-6800 (for maize) or LI-6400 (for rice) portable gas exchange systems (LI-COR Biosciences). The temperature, relative humidity, and photosynthetic photon flux density (PPFD) were set at 28 °C, 70%, and 1200 µmol m^-2^s^-1^ respectively for the leaf chamber. CO_2_ concentrations were set as 400, 200, 50, 100, 150, 200, 300, 400, 500, 600, 800, 1000, 1200 ppm in a step-wise manner. The A/Ci curve of maize was fitted according to Zhou *et al*., 2019.

### Blue native (BN)-PAGE and Western blot

Maize leaves were homogenized in cold STN medium (0.4 M sucrose, 50 mM Tris–HCl pH 7.6, 10 mM NaCl). The homogenate was filtered through two layers of nylon cloth and centrifuged at 200 g for 3 min at 4 °C. Supernatant was centrifuged at 5000 g for 10 min at 4°C to pellet crude chloroplasts. Chloroplasts were ruptured in cold TN medium (50 mM Tris–HCl pH 7.6, 10 mM NaCl), and thylakoid membranes were separated by centrifugation at 8000 g for 5 min at 4°C. The thylakoid membranes were suspended in solubilization buffer [25 mM BisTris-HCl, pH 7.0, 10 mM MgCl_2_, 20% (v/v) glycerol] at a final chlorophyll concentration of 1 mg ml^−1^. The thylakoid membranes containing the 0.5 mg ml^−1^ chlorophyll were solubilized with 1.2% (w/v) n-dodecyl-β-maltoside (DDM) by gentle agitation on ice for 1 h. After centrifugation at 15000 g for 10 min at 4°C, the samples were immediately subjected to Native-PAGE and loaded on chlorophyll basis of 5 µg per lane. Electrophoresis was performed at 4 °C by increasing the voltage gradually from 50 up to 200 V during the 5.5 h run. For two-dimensional analysis, excised BN-PAGE lanes were soaked in SDS sample buffer for 10 min, layered onto 12.5% SDS polyacrylamide gels, and electrophoresis was performed at 120 V.

For the immunoblot analysis of total proteins, leaves of 4-week-old plants were extracted in SDS sample buffer, and 25 µg of proteins were subjected to SDS-PAGE. The separated total proteins or proteins from two-dimensional gels were subsequently transferred onto polyvinylidene difluoride (PVDF) membranes (Merck, Cat. IPVH00010), hybridized with specific antibodies, and visualized by ECL assay kit (Thermo Scientific) according to the manufacturer’s protocol.

### Paraffin section and I_2_-KI staining

Leaf samples were harvested at 7AM and 7PM from 3-week-old plants. The middle portion of the fully expanded leaf was dissected and fixed in formaldehyde-acetic acid-alcohol (FAA) solution under vacuum (20 psi) at room temperature for at least 30min. Leaf sections were dehydrated using an ethanol series from 70% ethanol to 100% ethanol and incubated for 2 hours at each concentration solution. Samples were then infiltrated in varying concentrations of ethanol: histoclear solution (25% histoclear to 100% histoclear) and incubated for 2 hours at each concentration. Subsequently, samples were infiltrated in varying concentrations of histoclear: paraffin solution (25% paraffin to 100% paraffin) for 2 hours at each concentration. The processed samples were placed in heated 100 % paraffin and polymerized for 30 min at 4 °C. Embedded samples were cut into 10 μ m thick sections with a microtome (Leica RM2125RTS) and the paraffin sections were mounted on slides. Starch was visualized in leaf sections by iodine potassium iodide (I_2_-KI) staining.

### Histological staining of ROS accumulation

Leaf fragments of 2 cm size from the middle part of fully expanded leaves were transferred into DAB staining solution (1 mg ml^−1^ DAB, 10 mM Na_2_HPO_4_, and 0.05% Tween-20), vacuum infiltrated, and incubated for 6 h in the light (200 µmol photons m^−2^ s^−1^). Chlorophyll was removed by replacing DAB solution with decoloration solution (ethanol : acetic acid : glycerol = 3 : 1 : 1) and incubated at 85 °C. The samples were then processed with paraffin sectioning as described above, and cross sections were examined with light microscopy.

### Transmission electron microscopy

Leaf sections of 2 mm size were cut from the middle part of recent fully expanded leaves and fixed in 2.5% glutaraldehyde. The samples were placed in a microcentrifuge tube and low vacuum condition was applied for 60 min at room temperature. Post fixation in osmium tetroxide, embedding in Spurr’s resin, and other steps were performed by the technology platform in Center for Excellence in Molecular Plant Sciences, following standard procedures. The ultra-thin sections were imaged at 80 KV with a Hitachi H-7650 transmission electron microscope.

### RNA-sequencing

For maize, leaves at the youngest fully expanded stage were sampled, from 30-d-old plants. And for rice, samples were harvested from 60-d-old plants. Each biological replicate consisted of leaves from 4 individual plants, and 3 biological replicates (from 12 plants in total) were sequenced for each sample. Library preparation followed standard procedure, and the libraries were sequenced on Illumina Novaseq 6000 platform using 150-bp paired-end sequencing strategy. The cleaned reads of maize samples were mapped onto the Zm-B73-REFERENCE-NAM-5.0 reference genome, and cleaned reads of rice samples were mapped onto the RAP Gene ID reference genome, both using HISAT2. The differentially expressed genes (DEGs) were identified by DEseq2 (v1.16.1) with definition of fold change more than 2 and false discovery rate (FDR) <0.01.

### Gene expression analysis

Leaf samples were frozen with liquid nitrogen and ground in a tissue grinding machine. RNA was extracted using RNAiso Plus reagent (Takara, Cat. 9108) according to the manufacturer’s instructions. To synthesize cDNA, 2 µg RNA was used, with a first strand cDNA synthesis kit (YEASEN, 11141ES60). qRT– PCR was carried out with Hieff UNICON® Universal Blue qPCR SYBR Master Mix (YEASEN, 11184ES08) in a final reaction volume of 20 μL. Relative expression was calculated by comparison to *ZmEF1α* (Hughes and Langdale, 2020) or *OsACTIN*, and data processed using Excel software.

### Separation of BS strands

BS strands were isolated using a method modified from Chang *et al*., 2012. Maize leaves were harvested at the same stage for RNA-seq and 5 g of each sample were used to isolate BS strands. Each biological replicate consisted of leaves from 6 individual plants, and 3 biological replicates were used for each sample. Several leaf blades were stacked and cut perpendicularly to the long axis into 0.5- to 1-mm slices with sterilized razor blades and quickly transferred into 50 mL of cold BS cell isolation medium [0.6 M sorbitol, 50 mM Tris-HCl (pH 8.0), 5 mM EDTA, 0.5% polyvinylpyrrolidone-10, 10 mM DTT, and 100 mM 2-mercaptoethanol]. After blending for 10 s twice in a blender at high speed, the mixture was first filtered through a 178 µm mesh and then filter through a 125 µm mesh. The filtration processes were repeated. BS strands that passed through the 125 µm mesh were filtered and retained on a 70 µm mesh, then placed on a paper towel stack to remove excess moisture, and frozen in liquid nitrogen. The purity of BS strands was assessed with a light microscope and M and BS markers (PPDK and Rubisco, respectively) were monitored by Western blot **(Fig. S7c)**.

### Mass spectrometry analysis of protein samples

Equal amount of proteins (20 μg) were used for following analysis. Briefly, protein was reduced by 10 mM of TCEP and alkylated by 25 mM of CAA at 37°C in the dark for 1 h. Six volumes of cold acetone were subsequently added. Samples were left to precipitate overnight at −20°C and centrifuged at 12000 rpm for 20 min at 4°C, and supernatants were removed. Resulting pellets were washed twice with 90% cold acetone and suspended in 20 µL of 50 mM NH_4_HCO_3_. Sequence grade modified trypsin (Promega, Madison, WI) was added at the ratio of 1:20 to digest the proteins at 37°C overnight. The peptide mixture was desalted by C18 ZipTip and dried in a vacuum concentrator at 4°C. For MS analysis, the peptides were dissolved in 0.1% formic acid. Peptides were resolved on an EASY-Spray PepMap RSLC C18 column (Thermo Scientific, 25 cm x 150 μm ID, 2 μm, 55°C) with the following gradient profile delivered at 300 nl min^-1^ by a Dionex RSLCnano chromatography system (Thermo Scientific): 97% solvent A (0.1% formic acid in water) to 8% solvent B (0.1% formic acid in 80% acetonitrile) over 2 min, 8% to 23% solvent B over 90 min, 23% to 40% solvent B over 15 min, then 40% to 100% solvent B over 7min and maintained 8 min. The mass spectrometer was a Q Exactive HF-X hybrid quadrupole-Orbitrap system (ThermoFisher Scientific, MA, USA). Raw Data of DIA were processed and analyzed by Spectronaut 16 (Biognosys AG, Switzerland) with default settings. Maize proteome UP000007305_4577 (UniProt) database was referenced. Proteins entered into subsequent analysis must fulfill the following criteria: three or more unique peptides mapped to the protein or two unique peptides mapped to the protein but with at least 30% coverage. The relative protein amounts were calculated and the fold changes in *zmndf6* mutant relative to the wild type were presented.

### Metabolite profiling

Maize and rice samples were harvested from the same stages of plants used for RNA-seq. Each biological replicate consisted of 4 leaf discs (diameter=8 mm) from one individual plant, and at least 6 biological replicates (6 plants from 3 different mutant lines) were prepared for measurement. For each biological replicate, 4 leaf discs (approximately 40 mg) were flash frozen (dropped into liquid nitrogen within 1 second after cutting) and ground in a tissue grinding machine, 30 Hz × 1 min, three times. The ground powder was mixed with 800 μL extraction buffer [methanol: chloroform = 7:3 (v/v), −20°C pre-cooled], shaken in the tissue grinding machine (30 Hz for 3 min), and the mixture was kept in −20°C for 3-4 h. The mixture was added with 560 μL cold distilled water, shaken in the tissue grinding machine (30 Hz for 5 min), and put back in −20°C for 10 min. The sample was centrifuged at 2200 g, for 10 min at 4°C, and the supernatant (about 800 μL) was transferred to a new tube and kept in −20°C. For a 2^nd^ extraction, 800 μL of 50% methanol was added to the pellet and the tube was carefully vortexed for 15 s before put back in −20°C. After 30 min, it was centrifuged again at 2200 g, for 10 min at 4°C, and the supernatant (about 800 μL) was combined with last extraction. The total supernatant (about 1.6 mL) was filtered by 0.22 μm filter, and 5 µL were injected for HPLC-MS/MS analysis or for quality control.

Luna NH2 column (3μm, 100mm x 2mm, Phenomenex co. Ltd, USA) was used in liquid chromatography, and the separation was conducted with a gradient set of solution A (20 mM Ammonium acetate in 5% acetonitrile solution, adjusted to pH 9.5 with ammonia water) and solution B (acetonitrile): 0∼1 min, 15% A and 85% B; 1∼8 min, 70% A and 30% B; 8∼22 min, 95% A and 5% B; 22∼25 min, 15% A and 85% B. Mass spectrometry analysis of the eluent was performed with QTRAP 6500+ (AB Sciex, co. Ltd, USA) in multi reaction monitoring (MRM) mode, parameters were referred from previous work (Tang *et al*., 2022), and data analysis was performed using Analyst®1.6 software. Metabolites detected were identified by aligning the retention time to features of standard samples previous processed on the same platform (Tang *et al*., 2022). The relative amounts of each metabolite in different samples were calculated by the chromatographic peak areas, and the changes of the metabolites relative to the averaged values of the wild type were presented. The data presentation was adopted from Li *et al*., 2020.

### Accession numbers

Accession numbers for the genes measured by qRT–PCR are as follows: *ZmCA*(Zm00001eb037940), *ZmPEPC*(Zm00001eb383680), *ZmNADP-MDH* (Zm00001eb038930),*ZmPPDK*(Zm00001eb287770),*ZmME*(Zm00001eb1214 70),*ZmRbcS*(Zm00001eb197410),*ZmRbcL1*(Zm00001eb440790),*ZmRbcL2*(Zm00001eb187470), and *OsActin*(Os03g0718100).

## Results

### Construction of NDF6 and NDHU subunit deficient mutants in maize and rice

Chloroplast NDH is structurally divided into five subcomplexes: A, B, M(membrane), L(lumen) and ED(electron donor) **(Fig. S1a)**. Subcomplex B is composed of five subunits (PnsB1-5), and is specific to chloroplast NDH (Shikanai, 2015; Ma *et al*., 2021). By data mining, we found that the expression of *NDF6* (gene assigned as *PnsB4* recently) and *NDHU* (NDHU belongs to subcomplex ED) in maize BS cells was higher than in M cells, and higher in maize leaves than in rice leaves (Li *et al*., 2010; Wang *et al*., 2014) **(Table S1; Fig. S1b)**. In addition, their up-regulation amplitude in maize leaves relative to rice leaves was much higher than that of *NDHN* and *NDHO*, which had been reported in maize deletion mutants (Peterson *et al*., 2016), while *PGR5* and *PGRL1* were not highly expressed in maize leaves compared to rice leaves **(Table S1; Fig. S1b)**. To dissect and compare the significance of NDH-mediated CET in C_4_ and C_3_ photosynthesis, we generated maize and rice loss of function mutants of NDF6 and NDHU subunits using CRISPR-Cas9 technology.

For maize *ndh* mutants, two sgRNAs each were designed for the near 5’-terminal sequences of the first exons of *ZmNDF6* and *ZmNDHU* genes to construct gene editing vectors. Seven *ZmNDF6* edited and 5 *ZmNDHU* edited maize plants were obtained, and materials with 49bp, 51bp and 54bp homozygous deletions (starting at 16bp after ATG) in *ZmNDF6* gene were named *zmndf6-1*, *zmndf6-2* and *zmndf6-3*, respectively. Two homozygous mutation types in *ZmNDHU*, *zmndhu-1* and *zmndhu-2* were identified **(Fig. S2a, c)**.

For *ndh* mutants in rice, a single sgRNA near 5’ end of the third exon of the *OsNDF6* gene and the first exon of the *OsNDHU* gene was designed respectively to construct gene editing vectors. Twelve *OsNDF6* edited rice plants were obtained, including 3 *osndf6* homozygous lines with 3bp or 10bp deletions or a single base insertion. They were named *osndf6-1*, *osndf6-2* and *osndf6-3*, respectively. Seven *OsNDHU* edited materials were sequenced but all T0 plants were found to be heterozygous. After segregation, we identified homozygous *osndhu* mutants with single A-base insertion or 163bp deletion and named them *osndhu-1* and *osndhu-2* respectively **(Fig. S2b, d)**.

### Retarded growth of *ndf6* and *ndhu* mutants in maize but not in rice

After 30 days culture of T1 plants in a phytotron, the growth of *zmndf6* and *zmndhu* mutants was retarded compared with that of the wild type KN5585 **(Fig. 1a)**. Leaf chlorophyll contents and plant height of *zmndf6* and *zmndhu* mutants were significantly lower than those of the wild type **(Fig. 1b)**. The yellowish leaf colour of T1 plants was more obvious after 30 days in field greenhouse than in the phytotron **(Fig. S3)**. Rice *osndf6* and *osndhu* mutants showed no considerable difference in growth, chlorophyll content, and plant height compared with their wild type ZH11 **(Fig. 1c, d)**.

**Fig. 1.**
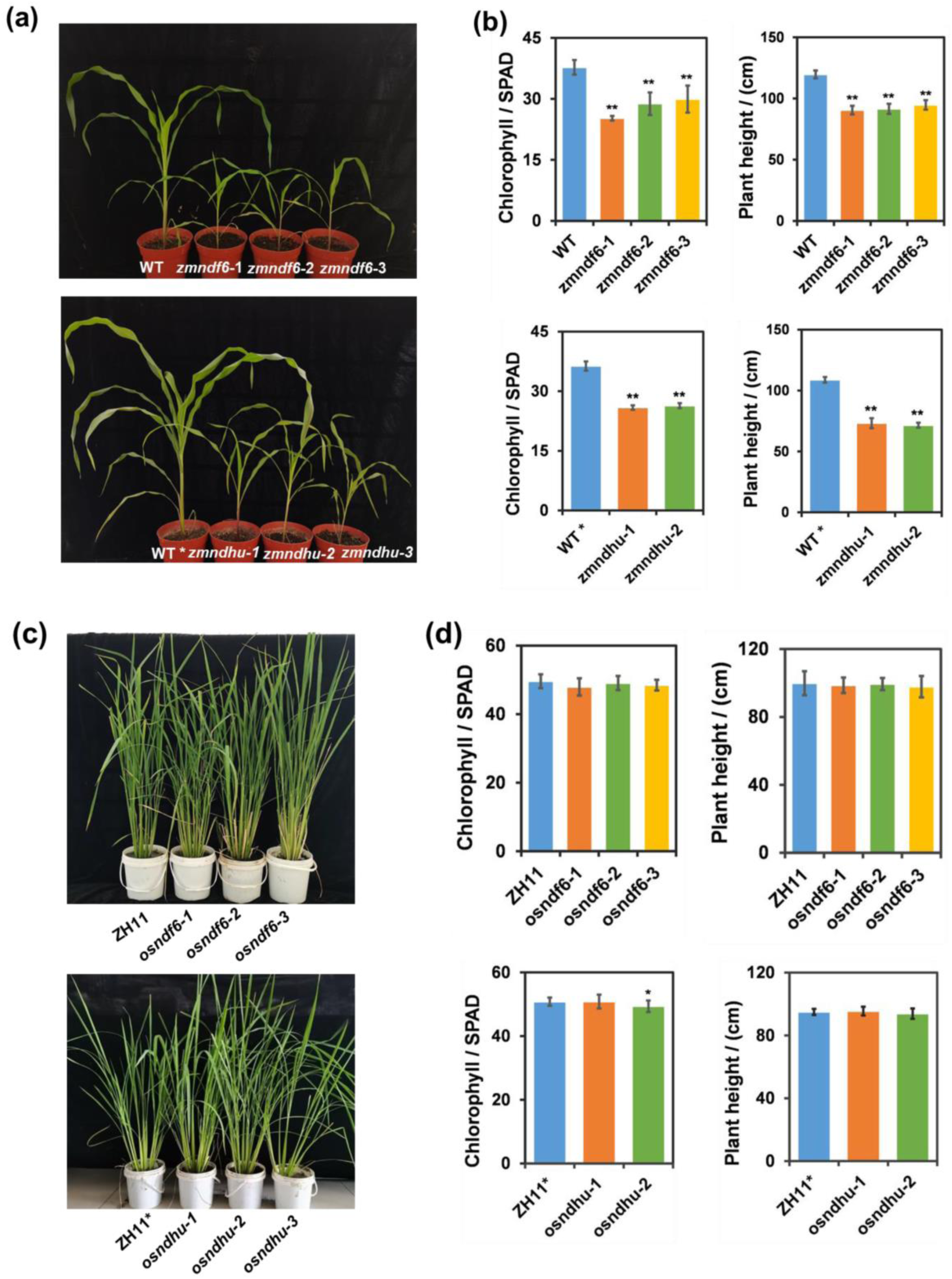
Altered growth phenotype in maize but not in rice *ndh* mutants **(a)** Maize plants of WT(KN5585) and *ndh* mutants 30 days after planting. **(b)** Comparisons of chlorophyll content and plant height between WT and *ndh* mutants. Data are mean ± SE (n = 6 biological replicates for chlorophyll content and plant height). **(c)** Rice plants of WT(ZH11) and *ndh* mutants 60 days after planting. **(d)** Comparisons of chlorophyll content and plant height between WT and *ndh* mutants. Data are mean ± SE (n = 12 and 8 biological replicates for chlorophyll content and plant height respectively). \**P* < 0.05, \*\**P* < 0.01 compared with WT according to Student’s *t* test. WT* indicates the wild type used in a different batch together with *zmndhu* mutants, and ZH11* indicates the wild type used in a different batch together with *osndhu* mutants, due to timing and availability of transgenic materials.

Post-illumination increase of chlorophyll fluorescence is used to indicate the operation of CET. When the driving force of LET via PSII is absent, NDH complex mediated electrons from the receptor side of PSI can flow back to reduce the inter-system PQ pool, causing a recovery in PSII chlorophyll fluorescence level (Asada *et al*., 1993; Mano *et al*., 1995; Mi *et al*., 1995). Compared with the wild type, the post-illumination increase of chlorophyll fluorescence disappeared in *zmndf6* and *zmndhu* mutants, demonstrating that the activity of CET was inhibited **(Fig. 2a)**. In addition, the decline of the chlorophyll fluorescence induction curve under actinic light was slower in the mutants than that in wild type, indicating a state of over-reduction of electron transport chain **(Fig. S4)**. Far-red light induces an increase in 810-830 nm absorption (representing oxidation of P700), and a decrease in light absorption after far-red light (representing reduction of P700^+^) reflects the rate of CET around PSI (Maxwell and Biggins, 1976; Mi *et al*., 1992; Asada *et al*., 1992). The dark reduction rate of P700^+^ in *zmndf6* and *zmndhu* mutants was significantly lower than that of the wild type **(Fig. 2b)**, indicating the partial inactivation of CET. Thus, a series of maize ZmNDF6 and ZmNDHU functional deletion mutants were successfully obtained.

**Fig. 2.**
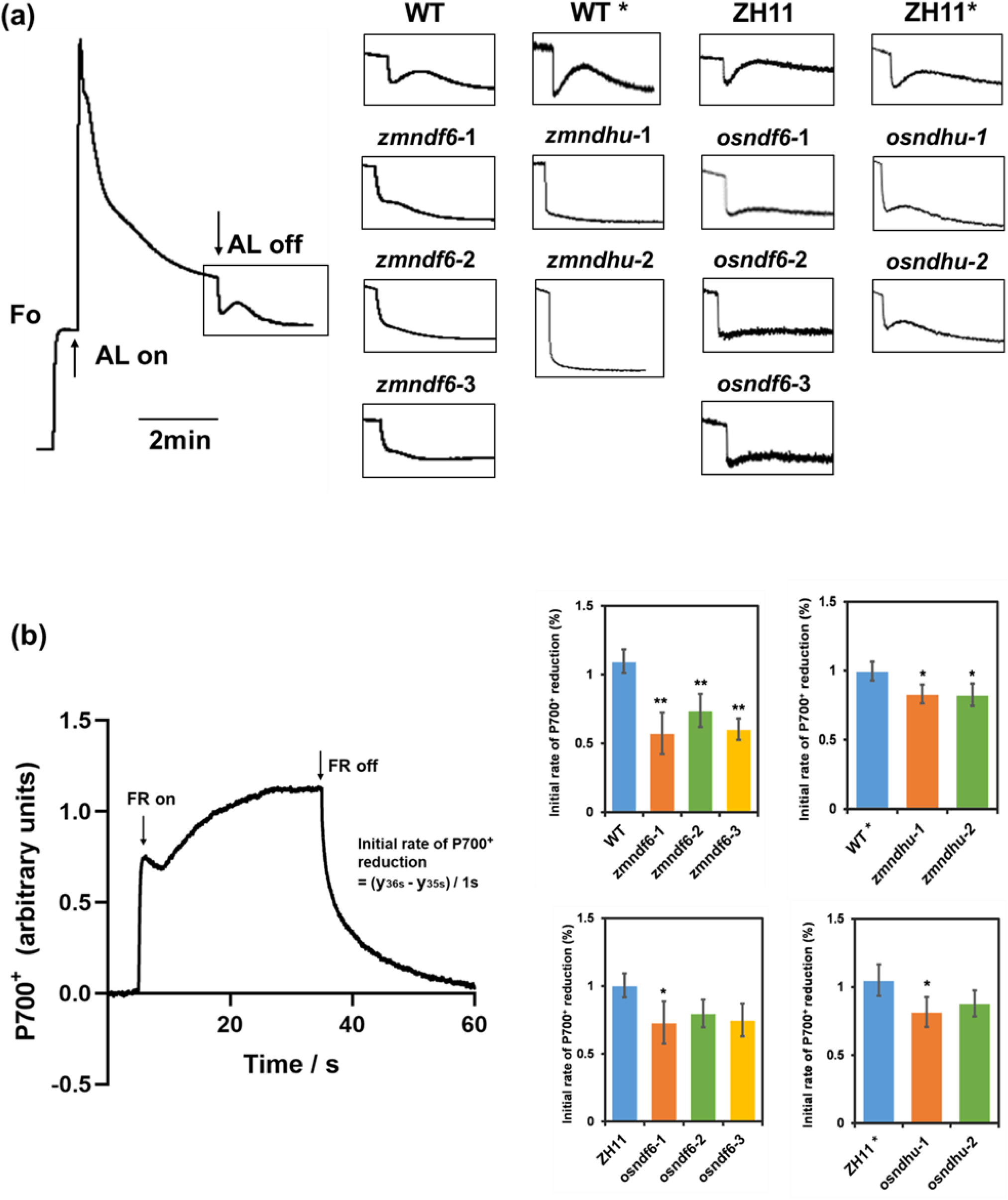
PSI cyclic electron flow was decreased in both maize and rice *ndh* mutants **(a)** Measurement of NDH cyclic electron transport activity, which was detected as a transient increase in Chl fluorescence (surrounded by a box) after turning off the AL. **(b)** The re-reduction of P700^+^ occurred after turning off the FR light, and comparison of the initial rates of P700^+^ re-reduction between WT and *ndh* mutants was calculated. Curves were normalized to the maximum P700 oxidation level. Data are mean ± SE (n = 4 biological replicates). \**P* < 0.05, \*\**P* < 0.01 compared with WT according to Student’s *t* test. WT* indicates the wild type used in a different batch together with *zmndhu* mutants, and ZH11* indicates the wild type used in a different batch together with *osndhu* mutants, due to timing and availability of transgenic materials.

To evaluate whether CET in rice NDH deficient mutants was affected, we measured the chlorophyll fluorescence and P700^+^ reduction in *osndf6* and *osndhu* leaves. Compared with the wild type, the post-illumination increase of chlorophyll fluorescence was decreased in *osndf6* and *osndhu* mutants **(Fig. 2a)**, but unlike in maize, there was no major difference in the descending phase of chlorophyll fluorescence under actinic light. The P700^+^ reduction rate of *osndf6* and *osndhu* mutants decreased to different degrees compared with that of the wild type **(Fig. 2b)**. These results show that the activity of the CET pathway was also impeded in the rice NDH deficient mutants.

### Photosynthetic CO_2_ assimilation was severely impaired in maize but not in rice NDH deficient mutants

To further explore whether and to what extent the photosynthetic activity of NDH deficient C_4_ and C_3_ plants was affected, we tested the CO_2_ response curve (A/Ci), electron transfer rate (ETR), non-photochemical quenching (NPQ) and other photosynthetic parameters of the maize and rice mutants. It was found that the initial slopes of the A/Ci curve, as well as the level of steady state photosynthetic rate, were clearly lower in maize *zmndf6* and *zmndhu* mutants than those in wild type **(Fig. 3a, b)**. The levels of ETR and NPQ in *zmndf6* and *zmndhu* mutants were also notably lower than those in wild type **(Fig. S5a, b)**.

**Fig. 3.**
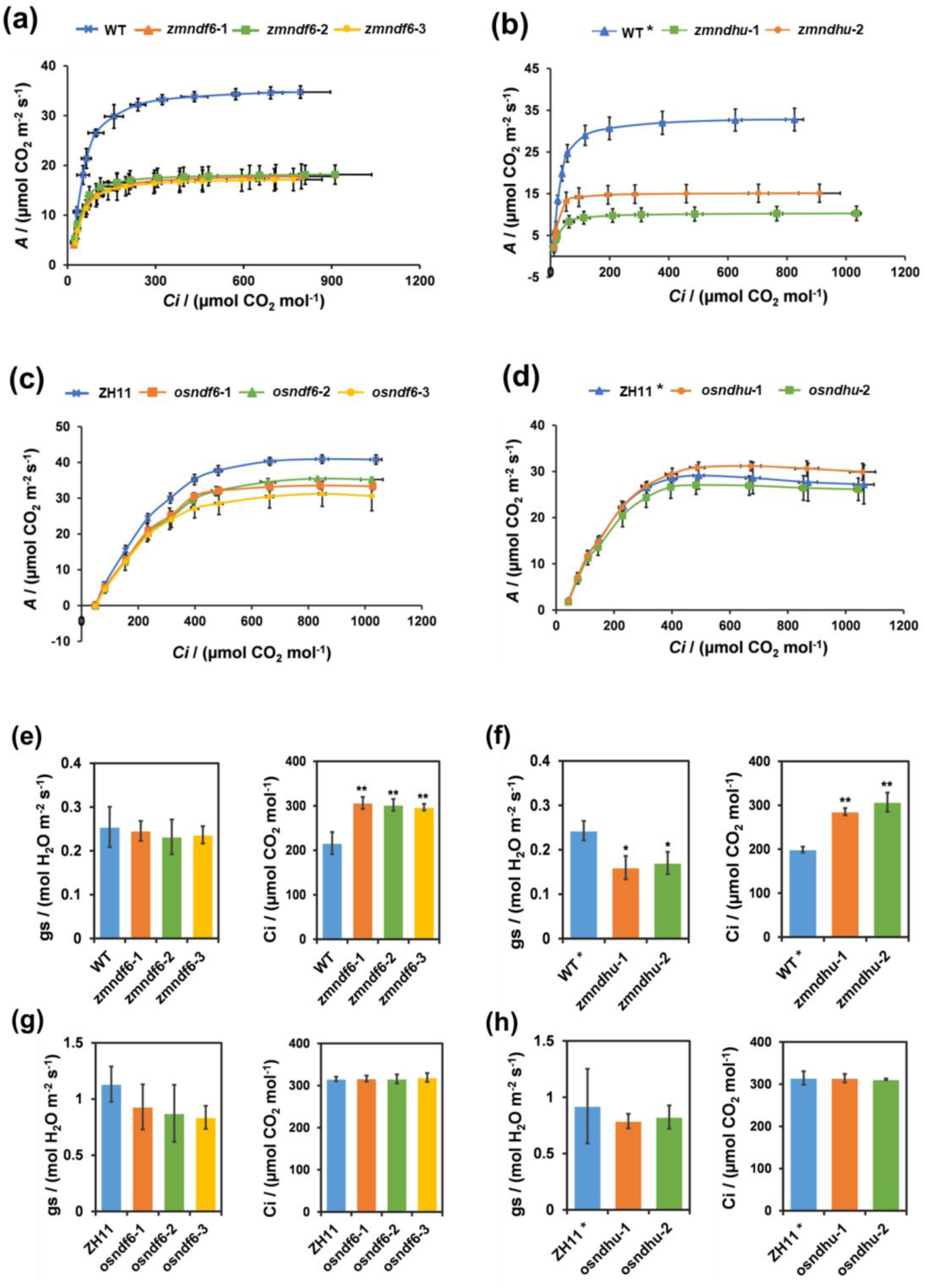
CO_2_ assimilation was severely inhibited in maize but not in rice *ndh* mutants **(a-d)** CO_2_ response curve of net photosynthetic assimilation rate (A/Ci) measured at 1200 µmol photons m^−2^ s^−1^ and 28 °C. (n = 3 biological replicates). **(e-h)** Stomatal conductance and intercellular CO_2_ concentration (Ci) measured at 28 °C and 400 µmol CO_2_ mol^-1^. Data are mean ± SE (n = 3 biological replicates). \**P* < 0.05, \*\**P* < 0.01 compared with WT according to Student’s *t* test. WT* indicates the wild type used in a different batch together with *zmndhu* mutants, and ZH11* indicates the wild type used in a different batch together with *osndhu* mutants, due to timing and availability of transgenic materials.

Different from the variation observed in maize *zmndf6* and *zmndhu* mutants, the initial slopes of the A/Ci curve, and the level of steady state photosynthetic rate, were not notably different in rice *osndf6* and *osndhu* mutants from those in wild type (except that the A/Ci curve of *osndf6* mutants appeared lower than wild type) **(Fig. 3c, d)**. Measurements of ETR and NPQ also showed no considerable difference between the rice *ndh* mutants and the wild type **(Fig. S5c, d)**.

In addition, it is noteworthy that the stomatal conductance (gs) of *zmndf6* and *zmndhu* mutants was not different or lower than that of the wild type, while intercellular CO_2_ concentration (Ci) was increased, especially in the 3 strains of *zmndf6* mutant **(Fig. 3e, f)**. This implies that CO_2_ assimilation was inhibited but it was not due to stomatal limitation. Changes of stomatal conductance and intercellular CO_2_ concentration in the rice *ndh* mutants were not obvious **(Fig. 3g, h)**.

### Decreased photosynthetic proteins and abnormal chloroplast ultrastructure in maize *ndf6* and *ndhu* mutants

To test whether the decreased photosynthesis of maize *ndh* mutants was related to decreased protein levels in the leaves, we compared the changes of protein components in PSI, PSII, cytochrome b6f, NDH, ATP synthase, and Rubisco between the mutants and the wild type. As expected, western blot analysis showed that NDF6 subunit was almost undetectable in the leaves of the *zmndf6* **(Fig. 4a)**, and NDHU subunit in *zmndhu* could barely be detected **(Fig. 4b)**. The protein levels of NDHH, NDHS, and NDHO subunits were also decreased in the maize *ndh* mutants **(Fig. 4a, b)**. The decrease of these subunits likely affected the amount and assembly of the NDH-PSI super complex as well, as reflected by blue-native gel and a 2nd-dimension electrophoresis **(Fig. 4e, f)**. By detecting the subunits of PSI and PSII, we found that the protein level of PsaA but not PsaD subunit tends to be lower in *zmndf6*, while PsaD but not PsaA subunit tends to be lower in *zmndhu* than in wild type. The protein level of PsbA (D1) was lower in both *zmndf6* and *zmndhu* than in wild type, while the PetA subunit of cytochrome b6f complex (PetA was found decreased in BS proteome, **Fig. S7a**) and the AtpB subunit of ATP synthase were not notably different from that of the wild type. Importantly, we found that the protein levels of RbcL in *zmndf6* (although RbcL is barely decreased in *zmndf6-1*) and *zmndhu* were notably decreased **(Fig. 4a, b)**, which supports the decreased CO_2_ assimilation activity of the maize NDH deficient mutants **(Fig. 3a, b)**.

**Fig. 4.**
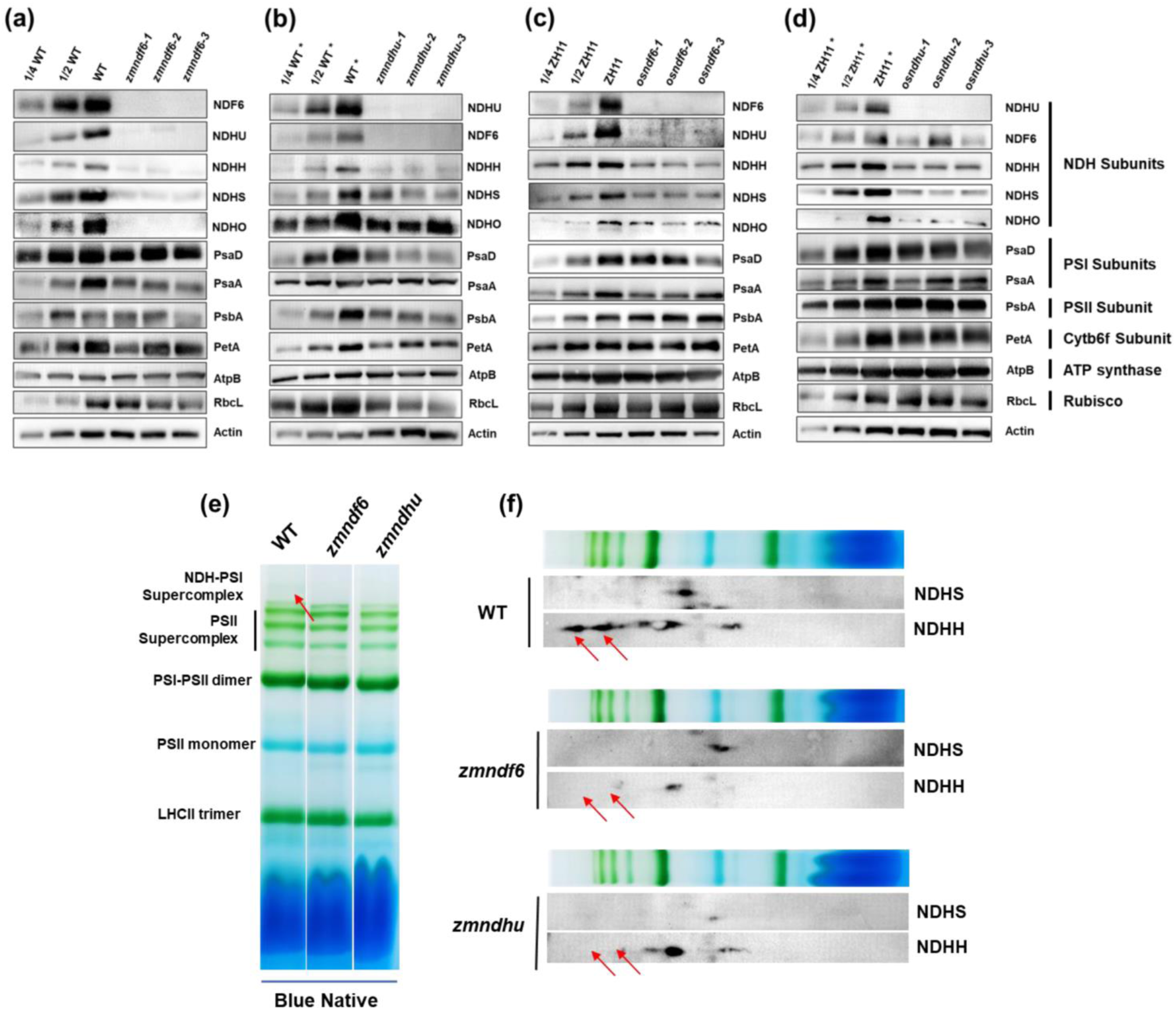
Immunoblot and BN-PAGE analysis reveals decreased amounts of photosynthetic proteins in maize *ndh* mutants The changes of selected protein components of PSI, PSII, NDH, Cytb6f, ATP synthase and Rubisco between WT and the mutants leaves. **(a)** Proteins in maize KN5585 and *zmndf6* mutants. **(b)** Proteins in maize KN5585 and *zmndhu* mutants. **(c)** Proteins in rice ZH11 and *osndf6* mutants. **(d)** Proteins in rice ZH11 and *osndhu* mutants. **(e)** Thylakoid membrane protein complexes isolated from WT, *zmndf6* and *zmndhu* were solubilized and separated by BN-PAGE (5-12%). High molecular weight green bands specific to WT are indicated. **(f)** Thylakoid membrane protein complexes separated by BN-PAGE were subjected to 2^nd^ dimension SDS-PAGE (12.5%), and the proteins were immunodetected with specific antibodies against NDHS, and NDHH, respectively. WT* indicates the maize wild type used in a different batch together with *zmndhu* mutants, and ZH11* indicates the rice wild type used in a different batch together with *osndhu* mutants, due to timing and availability of transgenic materials.

Protein levels of NDH subunits tested were also much decreased in the rice *osndf6* and *osndhu* mutants, but in contrast to the situation in the maize mutants, protein levels related to PSI, PSI, cytochrome b6f, ATP synthase, and Rubisco were not notably changed compared with the wild type (only PsaD and PsaA showed a slight downward trend) **(Fig. 4c, d)**. The differential changes in protein levels support the observation that photosynthetic electron transport, CO_2_ assimilation rates, and plant growth were compromised in maize but not rice *ndh* mutants, although CET activity was decreased in both.

To explore the relationship between the amount and integrity of NDH-PSI protein complex and thylakoid lamellar structure of maize BS chloroplasts, transmission electron microscopy was performed on maize wild type and *zmndf6* leaf samples. Compared with the wild type, non-stacking stromal thylakoid tended to be less assembled in the BS chloroplasts of maize *zmndf6* mutant, and areas with intermittent distribution or even absence of lamellae were observed **(Fig. 5a, b, e, f)**. For M cell chloroplasts, the thickness of the grana thylakoid was decreased and the grana lamellae were not well aligned in the *zmndf6* mutant compared with the wild type **(Fig. 5c, d, g, h)**. In addition, the starch granule accumulation of BS cells was lower in the mutant than in the wild type **(Fig. 5a, b, e, f)**. In accordance with this, starch staining by I_2_-KI in leaf paraffin sections showed that both the amount of starch synthesized early in the day and the accumulation late in the day were lower in the *ndh* mutants than that in the wild type **(Fig. 5j)**. The reduction in starch content is most probably caused by the decreased photosynthetic activity in maize *ndh* mutants **(Fig. 3)**.

**Fig. 5.**
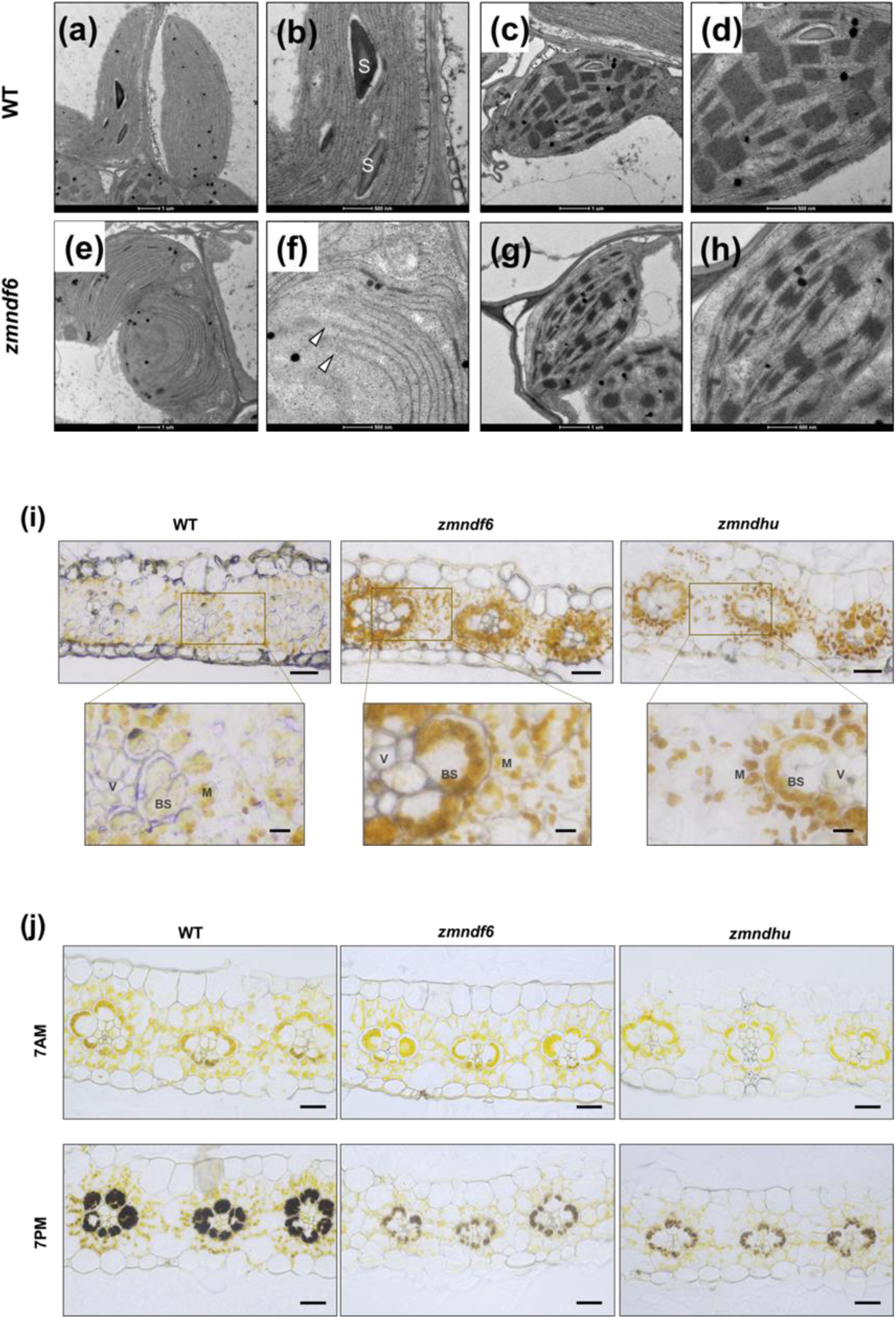
Abnormal chloroplast structure and accumulation of ROS and starch in maize *ndh* mutant leaves **(a-h)** Abnormal thylakoid ultrastructure of BS and M chloroplasts in maize *zmndf6* mutant. Transmission electron micrographs of BS chloroplast in WT **(a, b)**, M chloroplast in WT **(c, d)**, BS chloroplast in *zmndf6* **(e, f)**, and M chloroplast in *zmndf6* **(g, h)** are shown. “S” indicates starch granule. White triangles point to the broken ends of stromal thylakoid. **(i)** DAB stained transections showing over-production of ROS in the BS cells of maize *zmndf6* and *zmndhu* mutants. Brown signals were densely visible in the BS cells of *zmndf6* and *zmndhu* mutants, but not in those of the wild type. Lower panel images (scale bar = 5 μm) are enlarged from indicated area of the upper panel images (scale bar = 20 μm). **(j)** Light micrographs of transections showing less starch granules in maize *zmndf6* and *zmndhu* mutant leaves. Accumulation of starch grains in maize leaves at 7am and 7pm visualized by I_2_–KI (iodine in potassium iodide solution) staining. Scale bar = 20 μm.

### Up-regulated gene expression of Calvin-Benson cycle and cyclic electron transport in maize *ndh* mutants

In order to explore how and to what extent photosynthesis and other related pathways were affected at the transcriptional level, RNA sequencing was carried out on leaves of maize and rice NDH deficient mutants. We systematically analysed the differential expression of photosynthesis related functional genes, which were converted from a complete list of maize photosynthetic proteins demonstrated in Friso *et al*., 2010. Generally, gene expression in the two maize *ndh* mutants behaved differently from that in the rice *ndh* mutants, with transcript abundance linked with CET pathway and Calvin-Benson cycle mostly higher **(Fig. 6, for details with gene lists see Fig. S6)**.

**Fig. 6.**
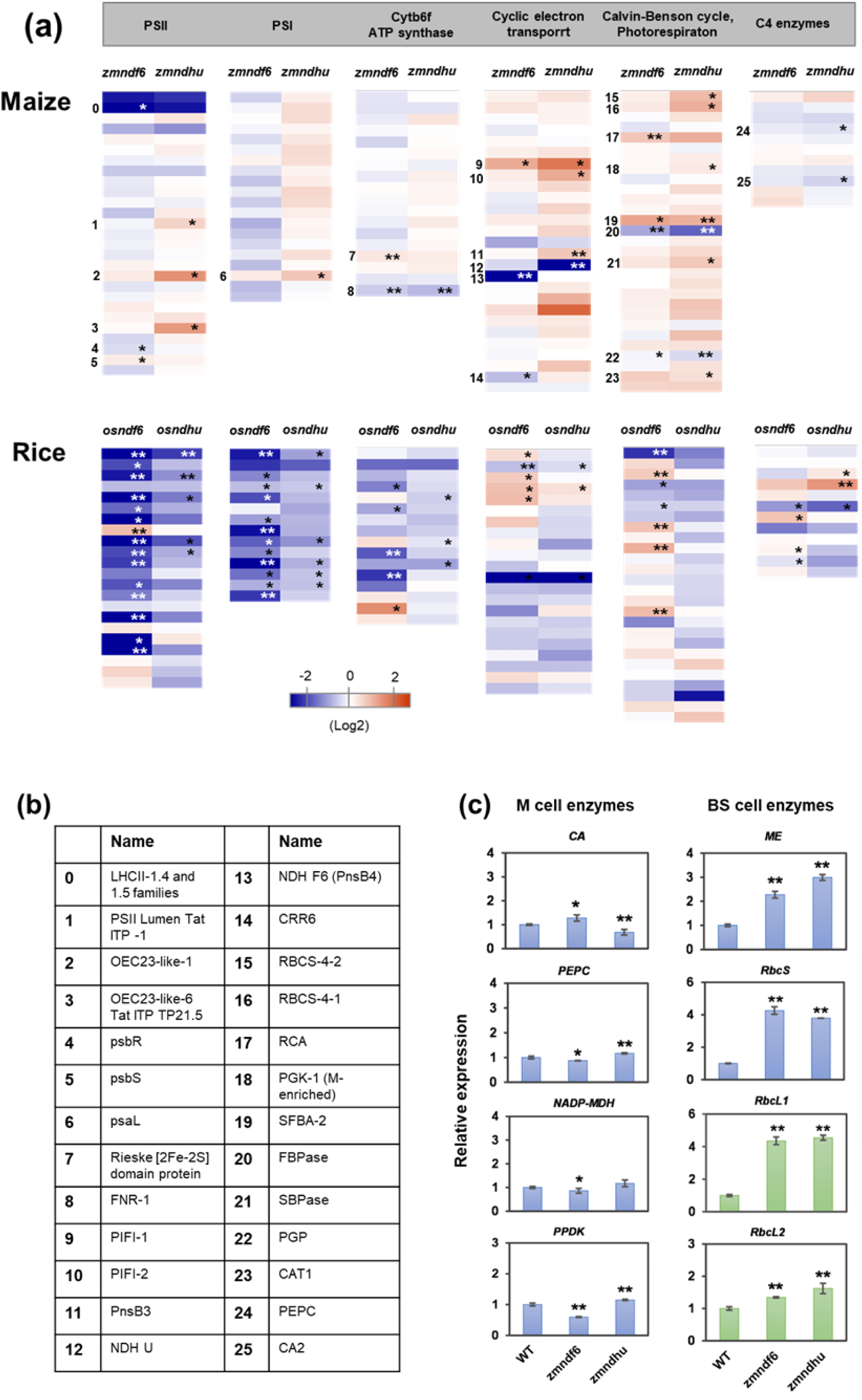
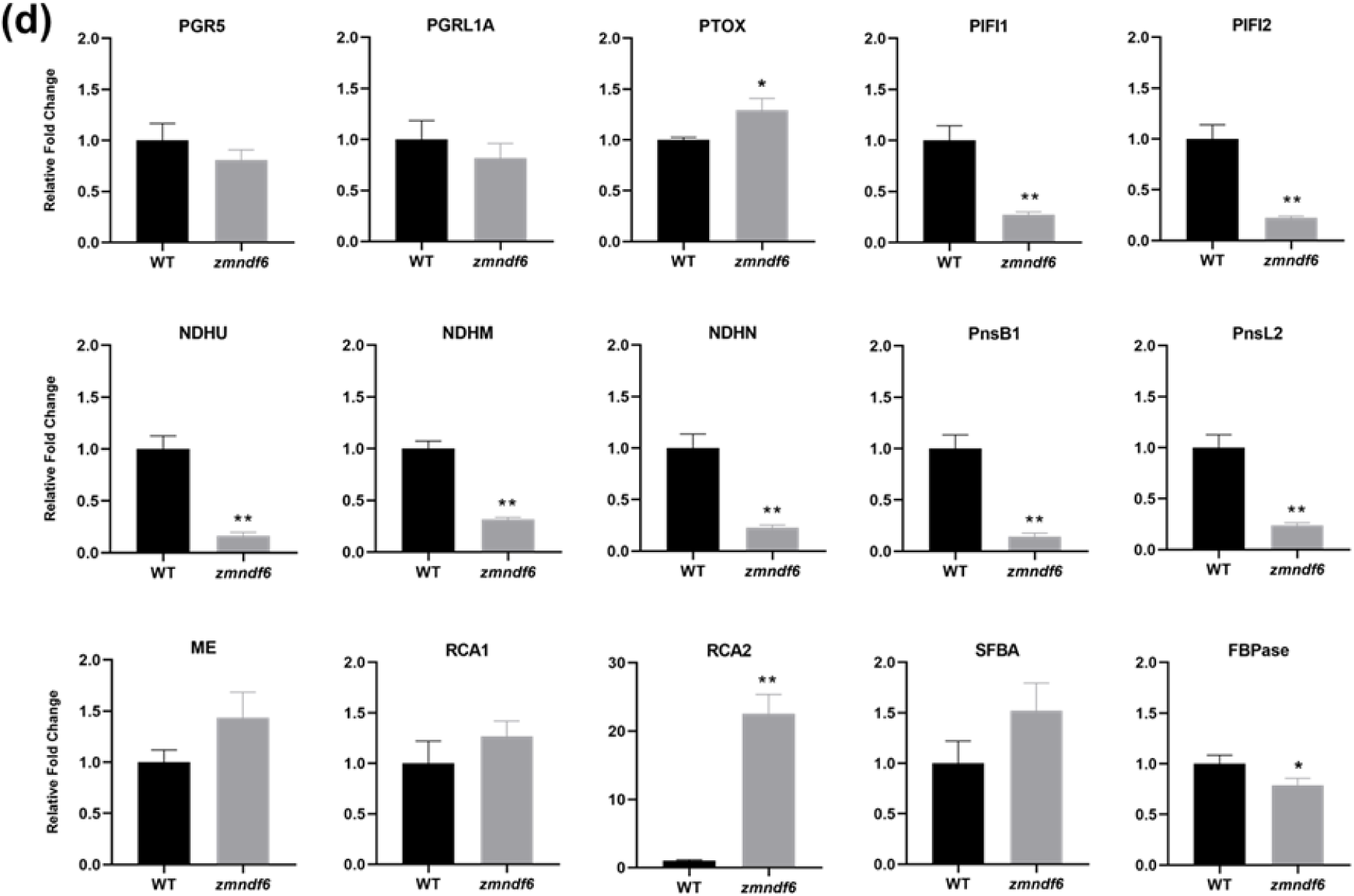
Changes of gene expression and BS cell protein content in maize *ndh* mutants **(a)** Heatmap shows differentially expressed genes in *ndh* mutants relative to their wild types (See **Fig. S6** for the heatmap with gene list). The significantly changed genes are assigned with numbers that link to names of encoded proteins in (b). The data represents mean value of 3 biological replicates. **(b)** Names of proteins encoded by significantly changed genes from (a). **(c)** RT-PCR validation showing increased transcript accumulation of BS but not M specific C_4_ genes in maize *ndh* mutants. *CA*, Carbonic anhydrase; *PEPC*, Phosphoenolpyruvate carboxylase; *NADP-MDH*, NADP malate dehydrogenase; *PPDK*, Pyruvate phosphate dikinase; *ME*, Malic enzyme; *RbcS*, Rubisco small subunit; *RbcL1*, Rubisco large subunit 1; *RbcL2*, Rubisco large subunit 2. **(d)** Changes of BS cell specific CET pathway and CO_2_ assimilation related protein contents. Comparisons were made between WT and *zmndf6* mutant according to quantitative values from BS cell proteomics. Data are mean ± SE (n = 3 biological replicates). \**P* < 0.05, \*\**P* < 0.01 compared with WT according to Student’s *t* test.

The expression of genes involved in LET, such as PSII, PSI, cytochrome b6f, and ATP synthase showed varied changes in the maize *zmndf6* and *zmndhu* mutants. Among this, the expression of genes encoding LHCII-1.4 and LHCII-1.5 were drastically and uniformly down-regulated (up to 8.6 fold) in the two maize mutants. Those up-regulated include genes encoding OEC23-like oxygen-evolving complex, PsbS, and Rieske domain protein **(Figs 6, S6a)**. As for the components related to CET, the *zmndf6* and *zmndhu* mutants displayed down-regulation of *NDF6* and *NDHU,* as expected, and lower transcript level of an extra gene of *NDF2*. Apart from that, the expression of PGR5 or other NDH-pathway associated nuclear genes showed consistent trends of up-regulation, including *PTOX* (plastid terminal oxidase) and *PIFI* (post-illumination chlorophyll fluorescence increase) **(Figs 6, S6b)**. In the rice *osndf6* and *osndhu* mutants, the expression of genes involved in LET showed consistent down-regulation. Interestingly, although most transcripts related to CET were down-regulated in the two rice mutants, *PTOX* and *PIFI* that are involved in chloroplast respiration were up-regulated **(Figs 6, S6a, b)**.

In maize *zmndf6* and *zmndhu* mutants, the expression of genes related to Calvin-Benson cycle and photorespiration was comprehensively up-regulated, especially for Rubisco activase (RCA) and Fructose-bisphosphate aldolase-2 (SFBA-2). One distinct exception is the markable down-regulation of FBPase in both maize *ndh* mutants **(Figs 6, S6c)**. Correspondingly, the expression of genes related to starch biosynthesis also showed extensive up-regulation **(Fig. S6d)**. In rice *osndf6* and *osndhu* mutants, there was no consistent up-regulation of Calvin-Benson cycle or photorespiration-related gene expression, indeed there was a tendency for down-regulation **(Fig. S6c)**. It should be noted that in line with increased levels Calvin-Benson cycle related transcripts in *osndf6* than in *osndhu* mutants, the expression of starch biosynthesis-related genes was also different between the two mutants **(Fig. S6d)**. This indicated the existence of intrinsic or physiological differences between rice *osndf6* and *osndhu* mutants.

### Effects of NDH deficiency on the BS specific C_4_ genes and ROS accumulation

Transcriptome analysis revealed that a deficient NDH pathway resulted in distinct changes in maize compared to rice at the level of gene expression. As chloroplast signals participate in nucleus-plastid communication (Surpin *et al*., 2002; Baier and Dietz, 2005), the redox changes within chloroplasts caused by lack of NDH **(Fig. S4)** may be one of the reasons for these gene expression changes. Specifically, we measured the gene expression of C_4_ enzymes in the maize *zmndf6* and *zmndhu* mutants by qRT-PCR. Consistent with the transcriptomic data, the expression of major genes (more abundant in transcripts) encoding BS cell localized ME, RbcS, and RbcL1 were up-regulated in *zmndf6* mutants, by 2.27, 4.26, and 4.35 fold, respectively. In contrast, the major gene expression of M cell localized PEPC, MDH, and PPDK did not markedly change or even decreased in the *zmndf6* mutants **(Fig. 6c)**. Similarly, an increased expression of BS specific C_4_ genes was observed in the *zmndhu* mutant.

To verify the redox changes in the chloroplasts of maize *ndh* mutants, we detected the accumulation of reactive oxygen species (ROS) in the BS and M cells by 3,3-Diaminobenzidine (DAB) staining **(Fig. 5i)**. DAB reacts with H_2_O_2_ to generate brown polymers that are stable in most solutions. With DAB uptaken by leaves, H_2_O_2_ can be localized *in situ* and at a subcellular level (Thordal-Christensen *et al*., 1997). Signal was densely exhibited in the BS chloroplasts of *zmndf6* and *zmndhu* mutants, and brown staining was also visible in the neighbouring M chloroplasts. In the wild type, overall staining intensity was much less than that in the *ndh* mutants **(Fig. 5i)**. These results indicate a remarkable increase in ROS accumulation in both BS and M cells when NDH is deficient in maize leaves, while in wild type leaves a balanced redox state contributes to lower ROS level. Less PSII and high NDH-CET is probably beneficial to prevention of ROS in the BS cells.

### Effects of NDH deficiency on the BS cell proteome

In order to further explore the effect of NDH deficiency specific in maize BS cells, we conducted a proteomic analysis by comparing the isolated BS cells of WT and *zmndf6* mutant. First of all, we found that contents of many NDH subunits (including NDHU, NDHM, NDHN, PnsB1, and PnsL2) significantly decreased, while contents of PGR5 and PGRL1 remained less changed, reassuring that the mutation of ZmNDF6 has reduced the abundance of NDH complex in maize BS cells **(Fig. 6d)**. The contents of BS specific C4 enzyme NADP-ME and Rubisco activase (especially RCA2) increased, although the contents of Rubisco subunits did not change (different from whole leaf Western blot, the similar amount of Rubisco here may be related to its highly enriched proportion in the BS of both WT and *zmndf6* mutant). Interestingly, consistent with the gene expression data, SFBA content enhanced whereas FBPase content declined. Further analysis of the proteins involved in starch synthesis revealed that beta-amylase (BAM3, BAM9), which is responsible for hydrolyzing starch to maltose, was significantly upregulated **(Fig. 6d; Fig. S7b)**.

On the other hand, proteins associated with PSI complex were significantly less in *zmndf6* mutant. Since PSI and NDH form super-complex, the deficiency of NDH complex in *zmndf6* mutant seems to also affect the abundance of PSI proteins. Components of ATP synthase appeared to be stable, while components related to Cytb6/f complex (PetA, PetD, and PetC for example) decreased. It is worth noting that STN7 and STN8 (involved in state transition), as well as FNR1 and FNR3 (responsible for electron transport between NADPH and Ferredoxin), were significantly up-regulated in *zmndf6* BS cells, indicative of remarkable changes in distribution of light energy and electron flow **(Fig. S7a)**. Analysis of the changes of proteins related to redox regulation showed significant downregulation of Thioredoxin and Peroxidase, indicated that loss of ZmNDF6 seriously affected the redox environment in maize BS cells **(Fig. S7b)**.

### Impaired NDH-CET caused critical changes in photosynthetic carbon metabolism in maize

The measured changes in photosynthesis, gene expression, and protein contents suggested that NDH deficiency had different impacts on CO_2_ assimilation in maize and rice. We therefore examined how NDH influenced photosynthetic metabolism using the maize and rice leaves for metabolic analysis. Metabolic profiles of the maize *zmndf6* and *zmndhu* mutants were found to be relatively consistent with each other, with obvious common up-regulated or down-regulated components, certifying the reliability of our biological samples and the effects caused by *NDH* mutations **(Fig. 7a)**. Specifically, metabolites involved in the Calvin-Benson cycle in maize *ndh* mutants were generally decreased compared with the wild type, except for RuBP and FBP, which accumulated significantly **(Fig. 7a, d)**. The accumulation of RuBP was consistent with the decrease in protein content of Rubisco large subunit **(Fig. 4a, b)** and the increase of intercellular CO_2_ concentration **(Fig. 3e, f)** in the mutants. In contrast to the decrease of most metabolites in the Calvin-Benson cycle, metabolites at earlier steps of photorespiration, such as 2PG, Glyco, Glx and Gly, increased in maize *ndh* mutants **(Fig. 7a, f)**, indicating an increase in photorespiration, which was consistent with a previous report (Peterson *et al*., 2016).

**Fig. 7.**
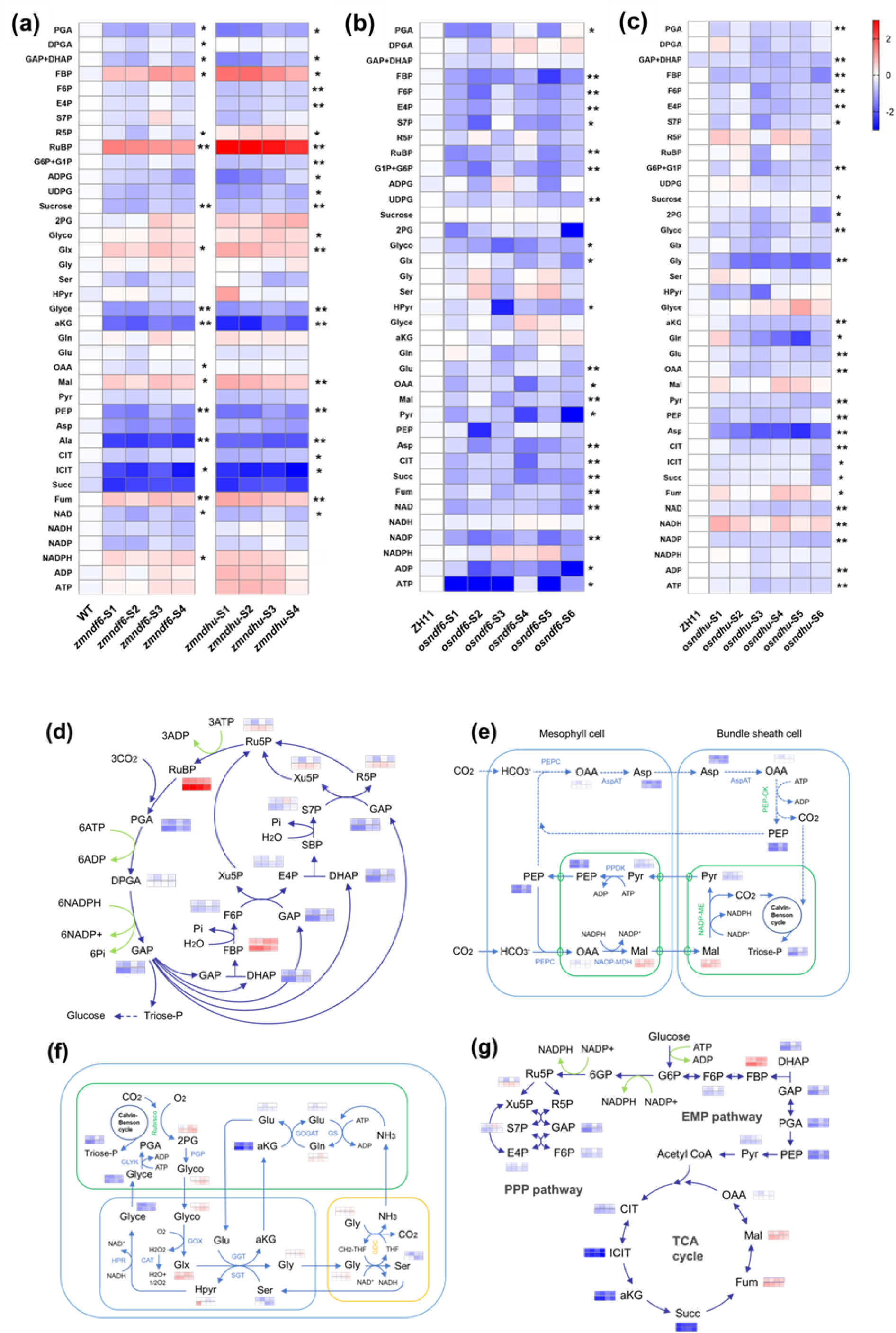
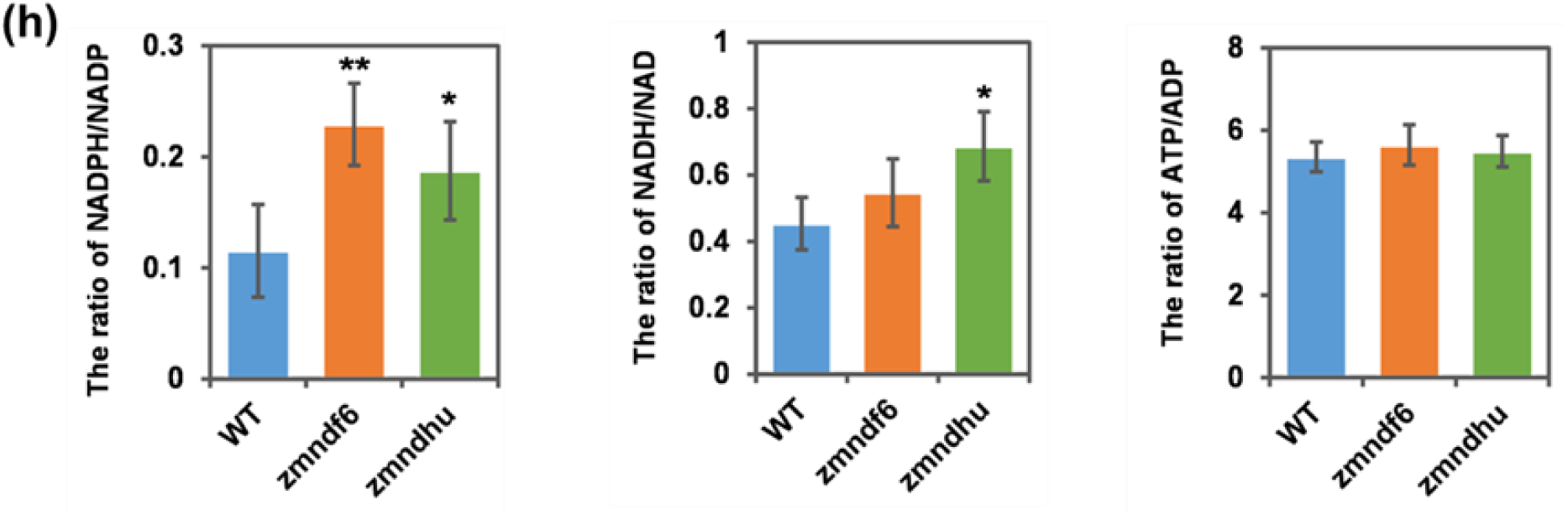
Loss of NDH-CET in maize leads to disturbance of key photosynthetic metabolites **(a)** Relative abundance (log2-transformed fold changes) of photosynthetic metabolites in maize *zmndf6* and *zmndhu* mutants. WT: The mean of log2-fold changes of 4 WT (KN5585) values relative to their average. S1-S4: Fold changes of mutant values relative to the average value of WT. **(b, c)** Relative abundance (log2-transformed fold changes) of photosynthetic metabolites in rice *osndf6* and *osndhu* mutants. ZH11: The mean of log2-fold changes of 6 WT (ZH11) values relative to their average. S1-S6: Fold changes of mutant values relative to the average value of WT. **(d-g)** Schematic diagrams showing key metabolites in **(d)** Calvin-Benson cycle (adapted from Figure 20-12, Biochemistry, sixth Edition, 2007, W.H. Freeman and Company), **(e)** C_4_ metabolic cycle (the dotted arrows pointing from PEP through OAA to Asp indicate the likely reduced activity of this pathway in the *ndh* mutants), **(f)** photorespiration pathway, and **(g)** EMP, PPP pathways, and TCA cycle of maize *ndh* mutants. Heatmap arrays derived from *zmndf6* (upper row) and *zmndhu* (lower row) are listed on the side of corresponding metabolites. EMP pathway, Embden-Meyerhof-Parnas pathway, also known as glycolysis. PPP pathway, pentose phosphate pathway. TCA cycle, tricarboxylic acid cycle. **(h)** Ratios of NADPH/NADP, NADH/NAD, and ATP/ADP. Data are mean ± SE (n = 3 biological replicates). \**P* < 0.05, \*\**P* < 0.01 compared with WT according to Student’s *t* test. Metabolite names showed in this figure are as follows: PGA (3-phosphoglyceric acid), DPGA (1,3-bisphosphoglyceric acid), FBP (Fructose 1,6-bisphosphate), GAP/DHAP(3-phosphoglyceraldehyde/Dihydroxyacetone), F6P (Fructose 6-phosphate), E4P (Erythrose 4-phosphate), S7P (Sedoheptulose 7-phosphate), PenP (Pentose phosphate, mixed of Ribose 5-phosphate, Xylulose 5-phosphate and Ribulose 5-phosphate), RuBP (Ribulose 1,5-bisphosphate), G6P/G1P (Glucose 6-phosphate/Glucose 1-phosphate), ADPG (Adenosine diphosphate glucose), UDPG (Uracil-diphosphate glucose), Sucrose (Sucrose), 2PG (Phosphoglycolic acid), Glyco (Glycolic acid), Glx (Glyoxylic acid), Gly (Glycine), Ser (Serine), Hpyr (Hydroxypyruvic acid), Glyce (Glyceric acid), aKG (α-Ketoglutaric acid), Gln (Glutamine), Glu (Glutamate), OAA (Oxaloacetic acid), Mal (Malic acid), Pyr (Pyruvate), PEP (Phosphoenol pyruvate), Asp (Aspartate), Ala (Alanine), CIT (Citric acid), ICIT (Isocitric acid), Succ (Succinic acid), Fum (Fumaric acid), NAD (Nicotinamide adenine dinucleotide), NADH (Nicotinamide adenine dinucleotide reduced), NADP (Nicotinamide adenine dinucleotide phosphate), NADPH (Nicotinamide adenine dinucleotide phosphate reduced), ADP (Adenosine diphosphate), and ATP (Adenosine triphosphate).

Extending the viewpoint to C_4_ metabolic cycle, we found that PEP, Pyr and Asp decreased to varying degrees, whereas malate increased in both *zmndf6* and *zmndhu* mutants **(Fig. 7a, e)**, which correlates with the increased gene expression and protein content of NADP-ME **(Fig. 6)**. Changes in malate probably also led to the accumulation of Fum, which interconverts with it in the tricarboxylic acid cycle (TCA cycle), while the levels of its upstream metabolites, such as CIT, ICIT, αKG, and Succ, decreased **(Fig. 7g)**. In addition, we have calculated the ATP/ADP, NADH/NAD and NADPH/NADP ratios, and the results indicate that the ATP/ADP ratio may not change, whereas the NADPH/NADP ratio is strongly increased in the two *ndh* mutants of maize **(Fig. 7h)**. The ratios are important because several NADPH participated C_4_ reactions are reversible, including the NADP-MDH and NADP-ME mediated reactions in the M and BS chloroplasts respectively. It should be mentioned that many of the metabolites are compartmentalized with a sub-pool being involved in C_4_ metabolism, the case of malate for example. It is possible that the changes are also among some other pool and not only in the pool that is involved in C_4_ metabolism.

The metabolic profiles of rice *osndf6* and *osndhu* mutants displayed more variable fluctuations, and an overall trend of decreased contents **(Fig. 7b, c)**. This metabolic variability may be related to the different gene expression patterns of the Calvin-Benson cycle between the two rice mutants **(Figs 6, S6c)**. On the other hand, while the levels of NADPH and adenylate in maize *ndh* mutants increased, the levels of them in rice *ndh* mutants appeared decreased **(Fig. 7b, c)**. These results support the remarkable difference in CO_2_ assimilation status observed between the maize and rice NDH deficient mutants.

## Discussion

To maintain an efficient Calvin-Benson cycle in the framework of the C_4_ pathway, it is necessary to dynamically balance the metabolism of both substance and energy. In this study, we show that NDH-mediated CET plays a crucial role in this process. The necessity and continuity of this regulation may be more prominent in C_4_ BS cells of maize, which provide a relatively special redox environment due to imported malate and NADPH. **Fig. 8** provides a summarized interpretation.

**Fig. 8.**
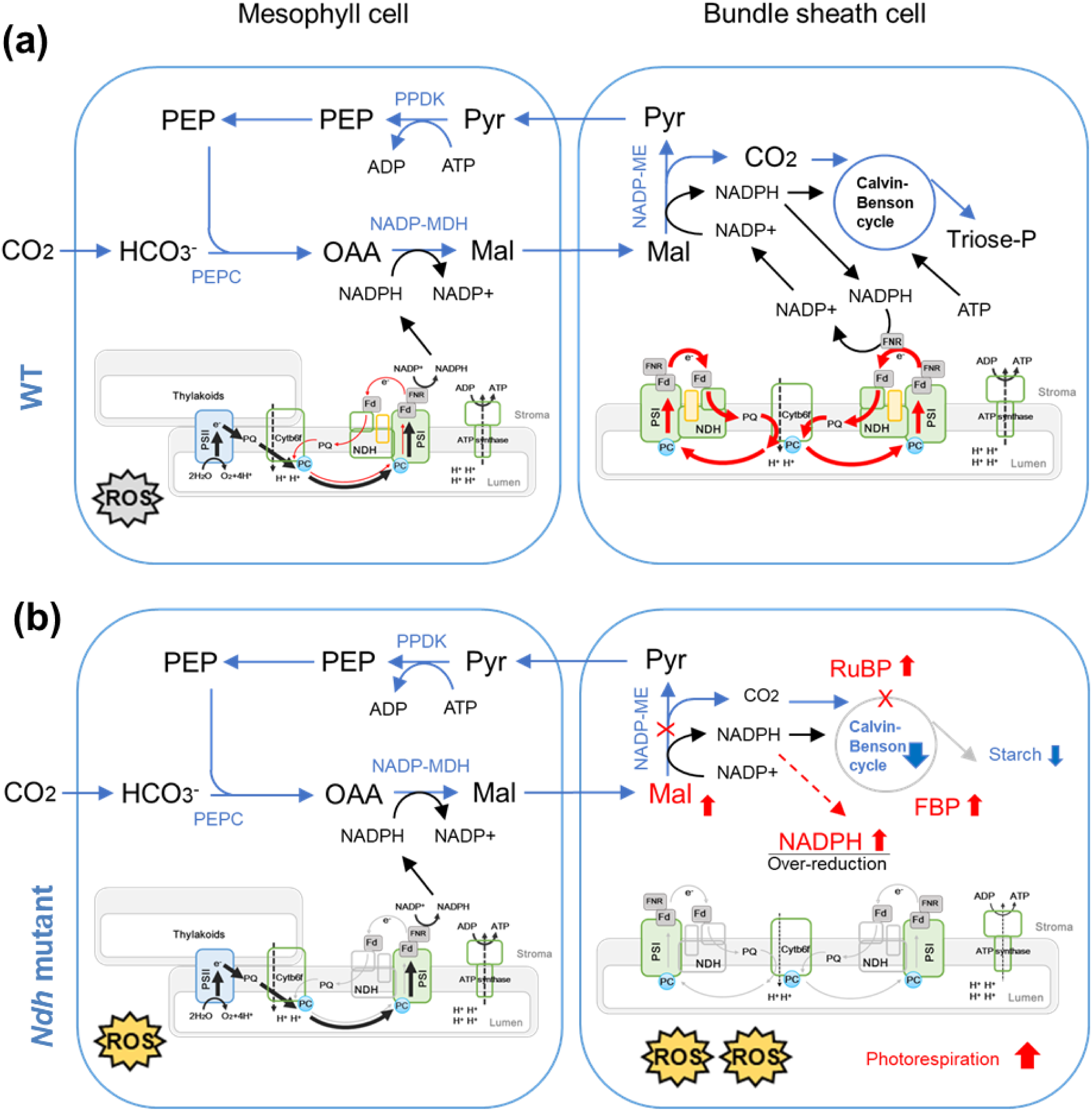
Schematic summary of the significance of NDH mediated cyclic electron transport in maize C_4_ photosynthesis (a) In the BS cells of wild type maize, malate is imported from M cells and mainly decarboxylated by NADP-ME, to release CO_2_ and couple NADPH generation. CO_2_ and NADPH is used in the Calvin-Benson cycle. Some of the NADPH may be involved in maintaining PSI-CET through the enriched NDH complex (bold red arrows compared to the thin red arrows in M cells), thus forming electron flows from malate to NADPH, from NADPH to PQ (via FNR, Fd, and NDH), and from PQ to PSI (through PC) or H_2_O (through PTOX), as there is almost no PSII-LET in the BS cells. Note: Maize BS cells have low levels of PSII, but we are not showing BS PSII in this diagram to save place and emphasize the enrichment of PSI; the electron flow from NADPH via FNR to Fd is reversible, while we have made the arrow single way for easier understanding; NDF6 (PnsB4) belongs to subcomplex B in orange colour. The process may be directly coupled with the CO_2_ pump, during which NADP^+^ is regenerated, ATP production is coupled, the input of malate along the concentration gradient continues, and Calvin-Benson cycle operates to fix CO_2_. It should be noted that 3PGA/triose-P shuttle is also important for delivery of ATP and NADPH to the BS cells, but is not presented due to tight schematic layout. ROS generation preferentially occurs in the M cells. (b) For a general comparison with wild type, in the BS cells of maize *ndh* mutants, PSI-CET is inhibited (thin grey arrows), with malate and NADPH accumulate (colored red), leading to redox imbalance, decreased CO_2_ fixation, increased ROS accumulation, and increased photorespiration. Since malate continues to be imported into maize BS cells and releases NADPH, the enriched NDH in BS cells functions to balance NADPH turnover, PSI electron flow, and Calvin-Benson cycle, apart from ATP supplement. Calvin-Benson cycle was inhibited in maize *ndh* mutants as reflected by the decrease of many metabolites but specific accumulation of RuBP and FBP. In addition, as combined effects of malate import and NDH deficiency, BS cells of maize *ndh* mutants suffered from an over-reduced condition. The altered redox status and metabolites flow should have disturbed the cellular environment, and casts effects on gene expression and protein accumulation patterns.

### NDH cyclic electron transport balances C_4_ metabolism and carbon flow in addition to ATP supply

Using NDH deficient mutants of maize and rice for comparison, we found that the metabolite and energy flows were effectively disrupted for C_4_ but not for C_3_ photosynthesis. Previous studies found that the direct electron donor of NDH was ferredoxin (Fd), and electrons were transferred between NADPH and Fd through ferridoxin-NADP^+^ reductase (FNR) (Ifuku *et al*., 2011; Peltier *et al*., 2016). Since NADPH is coupled to decarboxylation of the malate entering into BS cells, NDH may play a special role in maintaining the electron flow of PSI (the dominant component of BS chloroplasts), regeneration of NADP^+^, and the input of malate along the concentration gradient. This study demonstrates that both NADPH and malate tend to accumulate in maize *zmndf6* and *zmndhu* mutants, which is different from the situation in equivalent mutants in rice **(Fig. 7)**. The accumulation of malate and NADPH could be caused by the impaired NADPH turnover, and further caused by the decreased demand from Calvin-Benson cycle, as reflected by the accumulation of RuBP and lower levels of many other intermediates **(Fig. 7)** in maize *ndh* mutants. In contrast, photorespiration was favoured, leading to more production of 2-phosphoglyolate, and elevated levels of other intermediates in the first part of the pathway, such as glycolate and glyoxylate.

It is worth to note that the distinct down-regulation of gene expression for FBPase, together with the up-regulation of gene expression for fructose-bisphosphate aldolase (SFBA), echoes well with the FBP accumulation in maize *ndh* mutants **(Fig. S6c)**, indicating an important site specific regulation which is pending to be resolved. The accumulation of FBP also indicated an effect of the NDH pathway on rate-limiting steps of the Calvin-Benson cycle, and its involvement in the feedback regulation of CET, as the FBPase deficient mutant (*hcef1*) was reported to specifically upregulate NDH and increase CEF (Livingston *et al*., 2010a, 2010b). Embden-Meyerhof-Parnas pathway (EMP, also named glycolysis) is the primary step of cellular respiration and the universal pathway of glucose degradation. Pentose phosphate pathway (PPP) is an alternative route for carbohydrate degradation. As NADPH is generated in the G6P and 6GP dehydrogenization steps of the PPP pathway, and the PPP pathway connects to the EMP pathway via G6P **(Fig. 7g)**, whether the accumulation of FBP downstream of G6P in the EMP pathway was affected by NADPH accumulation is an open question, and FBP maybe also a hub for alternative carbon flow towards EMP and PPP pathway. Many EMP pathway and TCA cycle components decreased, with the exceptions of FBP, Fum, and Mal. Considering these pathways together, the contents of αKG and NADP^+^ decreased, while the contents of 2PG and RuBP increased in maize *ndh* mutants. Since the two metabolite pairs of “αKG, NADP^+^” and “2PG, RuBP”, play active or inhibitory roles in the CO_2_ concentration mechanism of cyanobacteria (Daley et al., 2012), it is possible that a similar regulatory role occurs in C_4_ plants.

On another hand, NDH-mediated CET is believed to be a key source of ATP required by C_4_ photosynthesis, especially in the BS cells of NADP-ME type C_4_ plants. This view is supported by the ability of NDH-mediated CET to couple additional ATP synthesis, the observed higher NDH content in C_4_ plants than in C_3_ plants, and the NDH enrichment in cell types requiring additional ATP for CO_2_ concentration (Takabayashi *et al*., 2005; Ishikawa *et al*., 2016b). The ATP/ADP ratio appeared not changed in *zmndf6* and *zmndhu mutants* compared with the wild type **(Fig. 7h)**. A possible explanation could be the decreased ATP consumption due to inhibited metabolic processes, although ATP supply was likely impaired in the maize *ndh* mutants.

### NDH cyclic electron transport poises C_4_ BS cell redox to regulate CO_2_ assimilation

The Calvin–Benson cycle is a redox regulated process. Many enzymes related to CO_2_ assimilation, including Glyceraldehyde-3-phosphate dehydrogenase (GAPDH), Phosphoribulokinase (PRK), Fructose-1,6-bisphosphatase (FBPase), Sedoheptulose-1,7-bisphosphatase (SBPase), NADP-dependent malatedehydrogenase (NADP-MDH), and Rubisco activase were known to be activated by the light through ferredoxin/thioredoxin-dependent reduction of regulatory disulfide bonds (Michelet et al., 2013). As BS chloroplasts have little PSII, disruption of NDH-CET may interfere with the provision of electrons for the reduction of ferredoxin and, hence, thioredoxin. The reduced thioredoxins are able to reduce regulatory disulfides and activate their target enzymes (Buchanan, 1991; Michelet et al., 2013). How and to what extent this is related to the interesting changes of Rubisco activase, FBPase, NADP-ME, beta-amylase and so forth in *zmndh* mutants, remain to be further examined. On the other hand, the accumulation of malate and NADPH in *zmndh* mutants is probably responsible for the over-reduced state of chloroplast stroma, and the obstructed status of electron transport chain **(Figs S4, S5)**. The increased ROS accumulation in the BS cells of *zmndf6* and *zmndhu* mutants **(Fig. 5i)** would be caused by the over-reduction, and this could negatively affect the content and activity of Rubisco, as well as Rubisco activase. The resultant inactivation of Calvin-Benson cycle in maize *ndh* mutants eventually leads to decrease of photosynthetic rate and the plant growth defects **(Figs 1, 3)**.

The decarboxylation of C_4_ acids in the BS cells is a key step in C_4_ photosynthesis. NADP-ME operation in the direction of decarboxylation is known to be hampered by high NADPH/NADP ratio, which has been recently highlighted by Andrea Brautigam (Brautigam et al., 2018), indicating that oxidized redox poise in the chloroplast stroma favors high velocity of decarboxylation. They summarized that BS chloroplasts achieve more oxidized state via the consumption of NADPH by Calvin-Benson cycle and the limited production of NADPH by abolishing PSII activity in BS cells. Here our results suggest a 3rd driving force of BS-enriched NDH-CET by NADPH turnover. In maize BS cells where malate is imported into and decarboxylated to release NADPH, it is possible that NDH-CET is needed to poise the NADPH level, so that its loss leads to the observed increase of the NADPH levels and the NADPH/NADP ratio, which in turn restricts decarboxylation by NADP-ME, leading to lower BS CO_2_ concentration, lower pyruvate content, and over accumulated malate level. The low BS CO_2_ concentration would restrict the Rubisco carboxylase reaction, together with the damage of Rubisco and Rubisco activase by accumulated ROS, explaining the high RuBP and lower levels of many other Calvin-Benson cycle intermediates.

As part of the early attempts to install C_4_ pathway, C_4_-specific NADP-ME was over-expressed in rice, but it led to bleaching of leaf color and growth hindrance, which resulted from enhanced photoinhibition due to an increase in the level of NADPH inside the chloroplast (Tsuchida et al., 2001). The malate and NADPH imbalance, pale green leaf, retarded growth, and ROS accumulation observed in our maize NDH function deficient mutants, are reminiscent of the effect of NADP-ME over-expression. The increased transcript level and BS protein content of NADP-ME strongly backup this scenario **(Fig. 6c, 6d)**. Therefore, a functional or enhanced operation of NDH-CET may be prerequisite when considering re-construction of the C_4_ cycle. In another hand, a recent work showed that high light stress led to more rapid ROS accumulation in BS than in M cells of rice, and also of C_4_ species (Xiong et al., 2021). The NDH deficiency resulted redox imbalance specific in maize BS cells should be responsible to their increased ROS level even under normal light.

### Deficiency of NDH disrupts the coordination of gene expression and protein accumulation in C_4_ plants

The compartmentalization of photosystems and C_4_ enzymes between BS and M cells is essential for the operation of C_4_ photosynthesis in maize. The regulatory mechanisms behind C_4_ differentiation remain unclear, but they probably involve the coordination of nuclear and plastid genomes, or specific changes in the intracellular environment to control gene expression in C_4_ leaves. Because of the abundance of the NDH complex in maize BS cells, and its important role discussed above, our maize *zmndf6* and *zmndhu* mutants can be viewed with cellular changes mainly in the BS. In particular, changes in the redox state of PQ pools in the chloroplast can affect nuclear gene expression through retrograde signal transduction (Foyer *et al*., 2012). Our data showed that the gene expression of the BS cell-specific C_4_ enzymes ME, RbcS, and RbcL1 were up-regulated remarkably, whereas the gene expression of M cell-specific enzymes PEPC, MDH and PPDK did not change or decreased in *zmndf6* mutants **(Fig. 6c)**. The BS specific protein contents of NADP-ME and Rubisco activase were consistently increased, but not the Rubisco subunits, which might be subject to more regulation or change on protein level.

With respect to cellular changes in M cells, Covshoff *et al*. identified a maize high chlorophyll fluorescence mutant (*hcf136*) with a PSII abnormality (Covshoff *et al*., 2008). Cell type-specific gene expression analysis showed that, compared with the wild type, the expression of M cell specific genes was promoted while that of BS cells was decreased. The authors suggested that the change of gene expression in M cells was related to retrograde signals from plastid to nucleus, while the change of gene expression in BS cells was more likely a secondary response caused by decreased reducing power or metabolite input (Covshoff *et al*., 2008). In another work, by studying the greening of *Cleome gynandra* leaves, combined with Norflurazon treatment, Burgess *et al*. pointed out that the regulation of C_4_ gene expression is considerably dependent on the chloroplasts, and light-induced activation of these genes is lost in damaged chloroplasts (Burgess *et al*., 2016).

Interestingly, according to the transcriptome analysis, apart from the CRISPR targeted *NDF6* and *NDHU* genes, the expression of many other NDH pathway-associated nuclear genes, as well as *PGR5* and *PTOX*, showed various degrees of up-regulation **(Fig. S6b)** in maize *zmndf6* and *zmndhu* mutants. One possible trigger could be the increased ROS accumulation in these mutants, as Strand *et al*. found that treatment of leaves with H_2_O_2_ could improve the NDH complex-dependent CET activity (Strand *et al*., 2015). It could be also partly attributed to the feedback signals, from the decreased protein levels of many NDH subunits. Equally remarkable, Calvin-Benson cycle related gene expression were comprehensively and consistently up-regulated in the two maize *ndh* mutants **(Fig. S6c)**. We suspect that these parallel changes of gene expression between CET and Calvin-Benson cycle are closely related or feedback affected by the changes of protein levels **(Figs 4, 6d)**, metabolic flows **(Fig. 7)**, or redox state. It could be also a projection of the intrinsic involvement of NDH pathway in the regulation of C_4_ carbon metabolism. Studies on the genus Flaveria indicated that NDH increased during the evolution of C_4_ pathway, especially during the transition stage from C_3_-C_4_ intermediate to C_4_-like (Nakamura *et al*., 2013), further suggesting that the function of NDH in NADP-ME C_4_ plants may be more closely related to the regulation of carbon metabolism. However, it remains difficult to identify genetic regulators that systematically upregulate NDH expression.

In conclusion, as modelled in **Fig. 8**, we propose that NDH cyclic electron transport forms an indispensable regulatory circuit for the two-celled C_4_ photosynthetic system in maize. The significance of the NDH pathway in C_4_ plants can be attributed to both ATP supply and NADPH turnover, as well as the adjustment of malate flux and NADP-ME activity. The loss of NDH function leads to metabolic, redox, and other regulatory imbalances, especially in BS cells, which then feedback to affect the coordination of photosynthetic gene expression and protein levels. Our study advances the functional understanding of NDH enrichment in maize BS cells, and hopefully will inspire fine tuning strategies to be re-considered towards engineering C_4_ photosynthesis.

## Supporting information

Dataset S1 Transcriptome data of maize zmndf6 and zmndhu mutants

Dataset S2 Transcriptome data of rice osndf6 and osndhu mutants

Dataset S3 Expression of photosynthetic genes in maize and rice ndh mutants-V2

Dataset S4 BS cell proteomics of maize WT and zmndf6 mutant

Dataset S5 Metabolite analysis of maize and rice ndh mutants-V2

Dataset S6 Summary of statistics-V2

Dataset S7 Summary of primers

## Acknowledgments

We thank Prof. Hualing Mi and Prof. Jane Langdale for critical comments on this work. We thank You Zhang and Luhuan Ye for helping with gas exchange, chlorophyll fluorescence, and P700 measurements; Zai Shi, Rui Zhang, and Yanfei Fan for other material and instrument assistance. We are grateful to Xiaoyan Gao, Zhiping Zhang, Jiqin Li, Wenli Hu, Shanshan Wang and Xiaoyan Xu for technical support on TEM and HPLC-MS; Chunguang Chen from ORIZYMES for antibody generation and Mi lab for gift of other antibodies; WIMI Biotechnology for maize and rice mutant generation. This research was funded by Strategic Priority Research Program (No. XDA24010203-2), the National Natural Science Foundation of China (No. 31970257), and a joint grant for international exchange from Royal Society and National Natural Science Foundation of China to Andrew J. Fleming and Peng Wang (No. 32011530166).

## Author contributions

QZ and PW conceived the project; QZ performed most experiments, data analyses and produced the figures; ST analyzed the RNA-seq data; QT and GC provided essential help for metabolite measurement. QZ, YZ, AJF, XZ and PW interpreted the results, wrote, and revised the paper with input from all authors.

## Data availability

The mRNA-seq datasets generated in this study have been deposited into the SRA database at NCBI under BioProject IDs PRJNA843342.

## Supporting Information

**Fig. S1.**
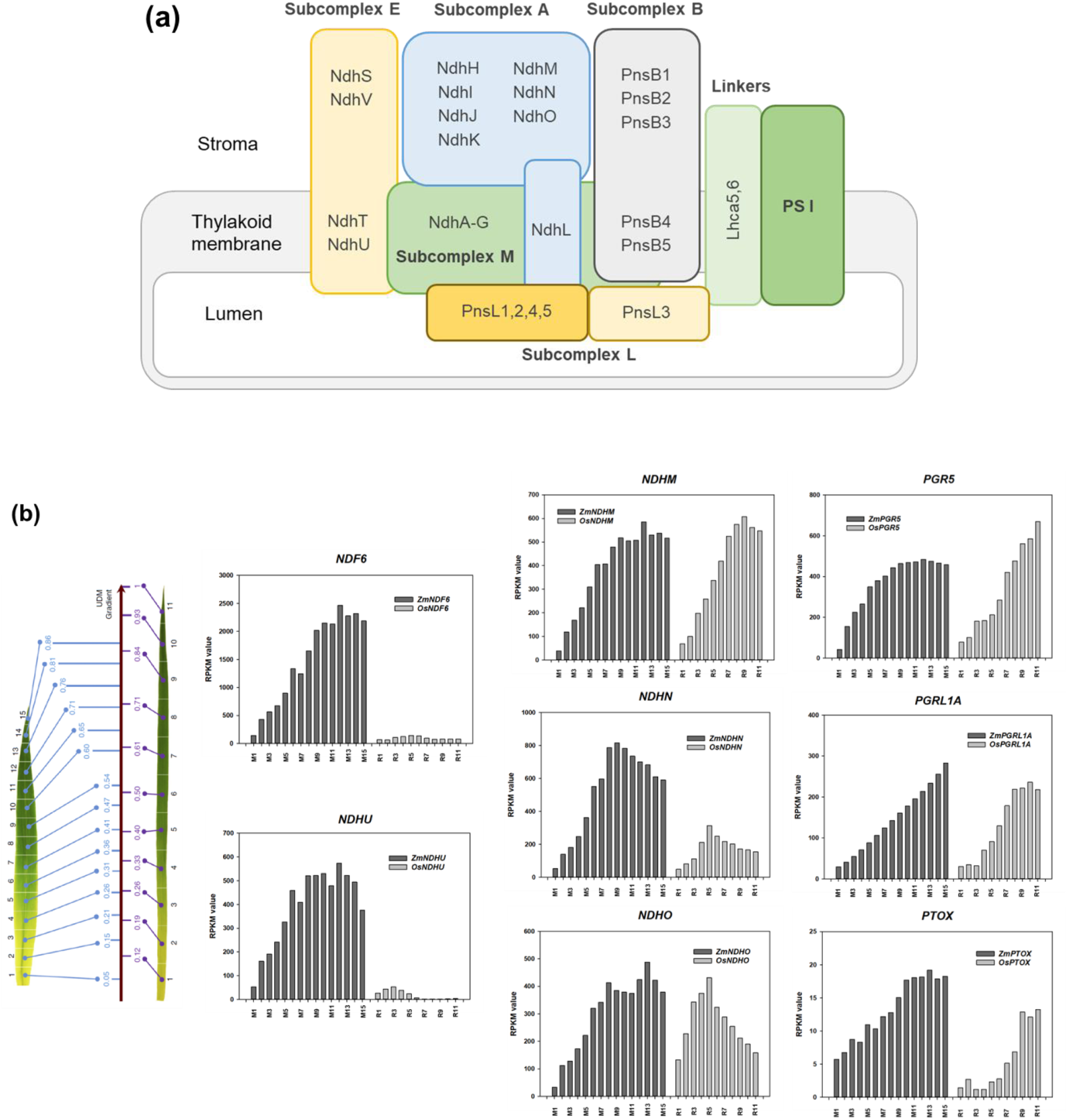
Comparison of the transcript levels of cyclic electron transport related components in maize and rice leaf gradients. Schematic model and transcript levels of NDH subunits in maize and rice (a) Schematic model of the NDH–PSI supercomplex (Derived from Shikanai, 2015). NDF6 was renamed as PnsB4; (b) Comparison of the transcript levels of cyclic electron transport related components in maize and rice leaf gradients (Derived from Wang et al., 2014).

**Fig. S2.**
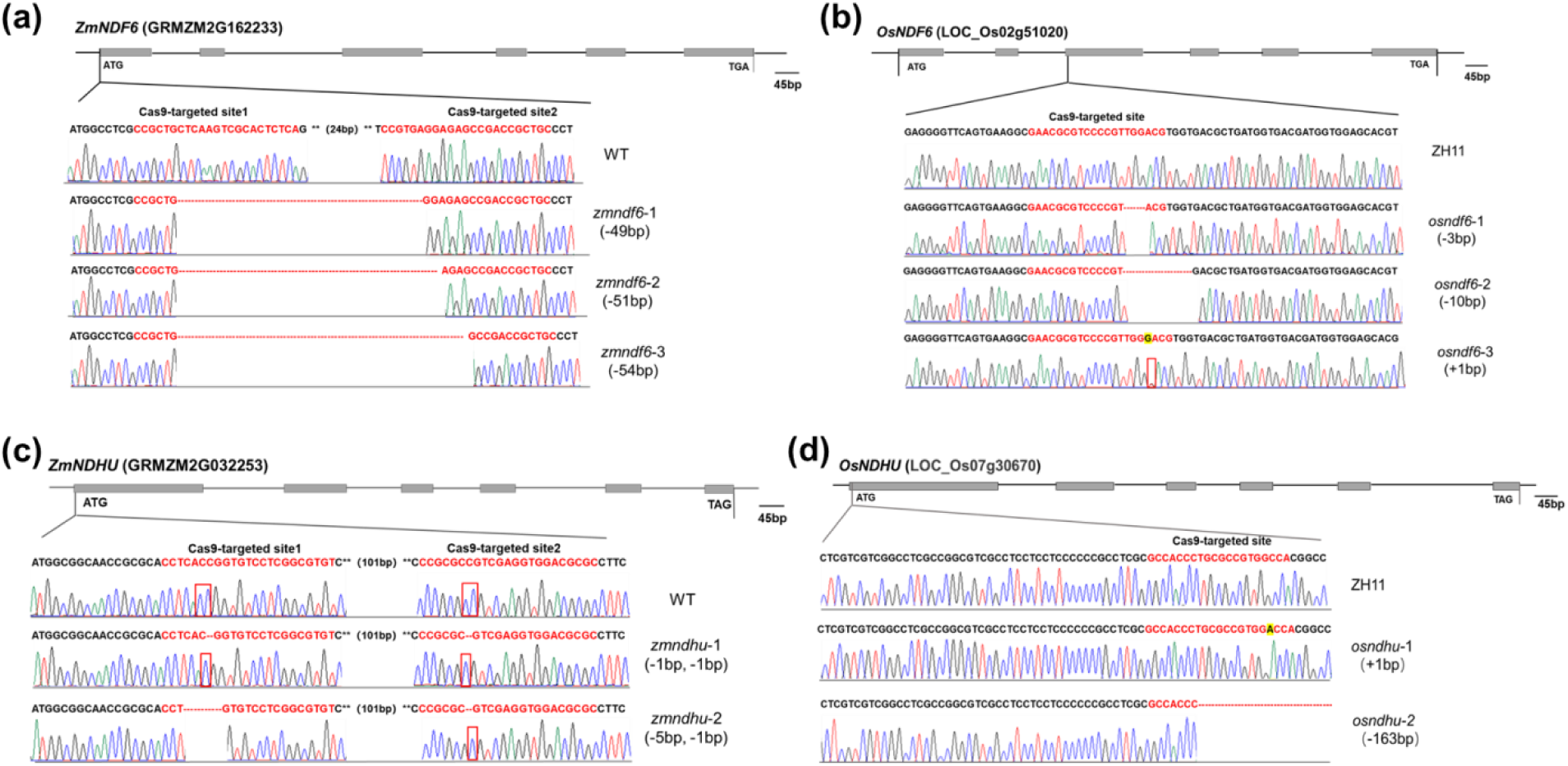
Mutation of NDH genes in maize and rice with CRISPR-Cas9 technology Schematic figures of the target sites and CRISPR-Cas9 edited sequences for (a) *ZmNDF6*, (b) *OsNDF6*, (c) *ZmNDHU*, and (d) *OsNDHU*. Nucleotides in red represent the Cas9 target site and red box indicates the difference between WT and mutants. See main text for details.

**Fig. S3.**
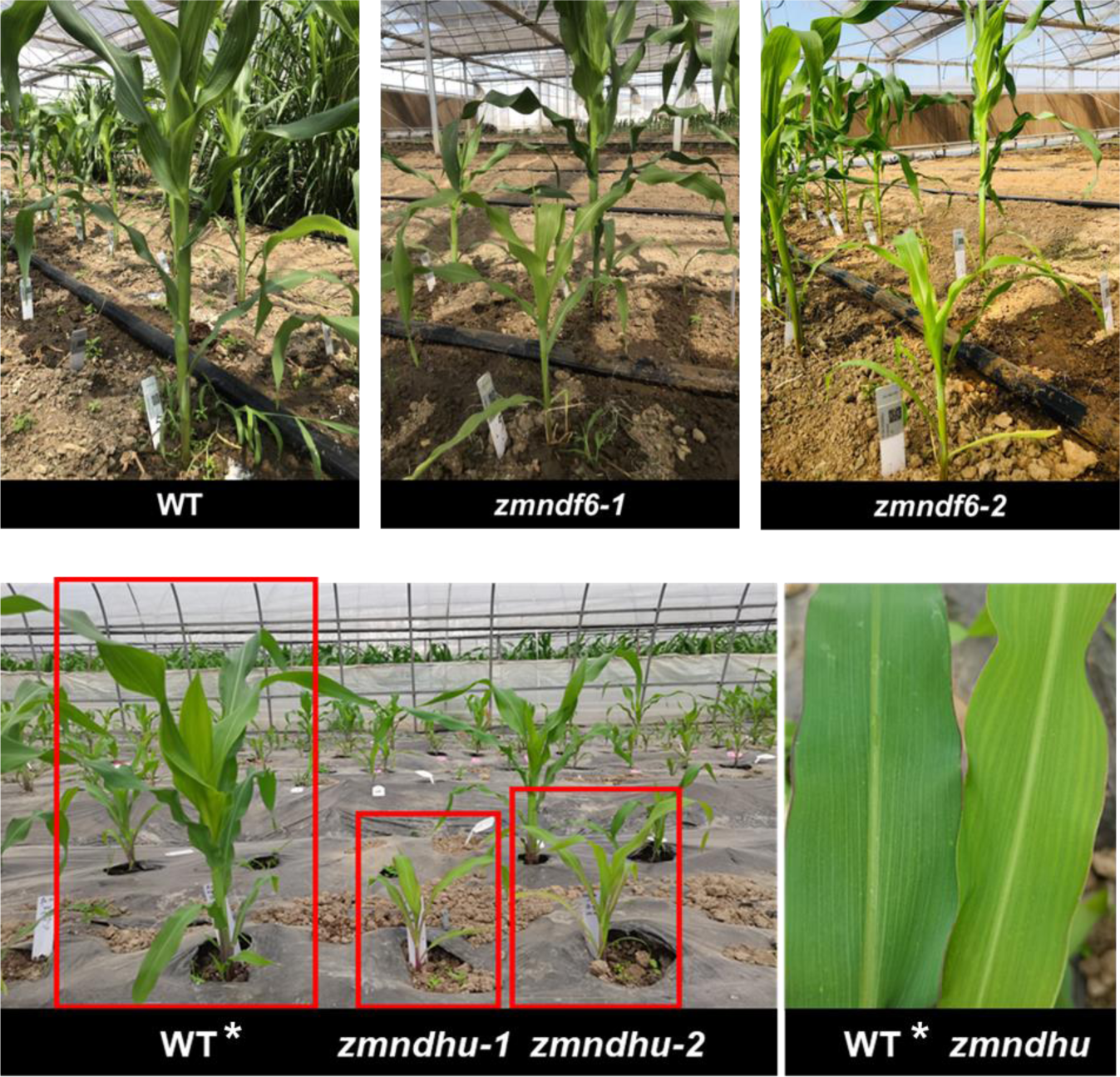
The phenotypes of maize *zmndf6* and *zmndhu* mutants growing in the field greenhouse

**Fig. S4.**
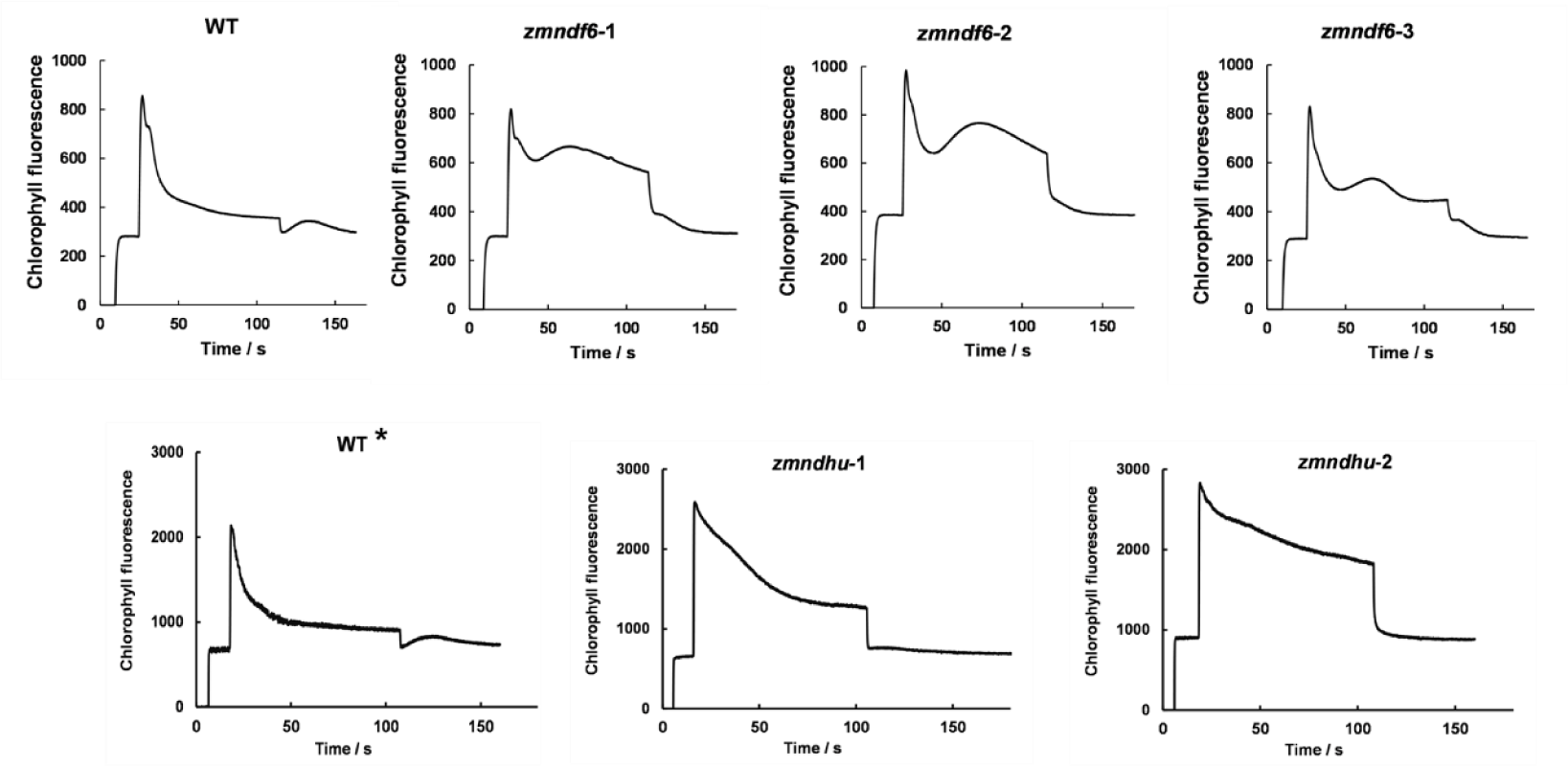
Induction Kinetics and Post-illumination Increase in Chlorophyll Fluorescence in Maize Plants The post-illumination increase of Chlorophyll fluorescence disappeared in *zmndf6* and *zmndhu* mutants, showing the inhibited activity of cyclic electron transport. The decline of Chlorophyll fluorescence under actinic light in the mutants was slower than that in the WT, indicating over-reduction of electron transport chain.

**Fig. S5.**
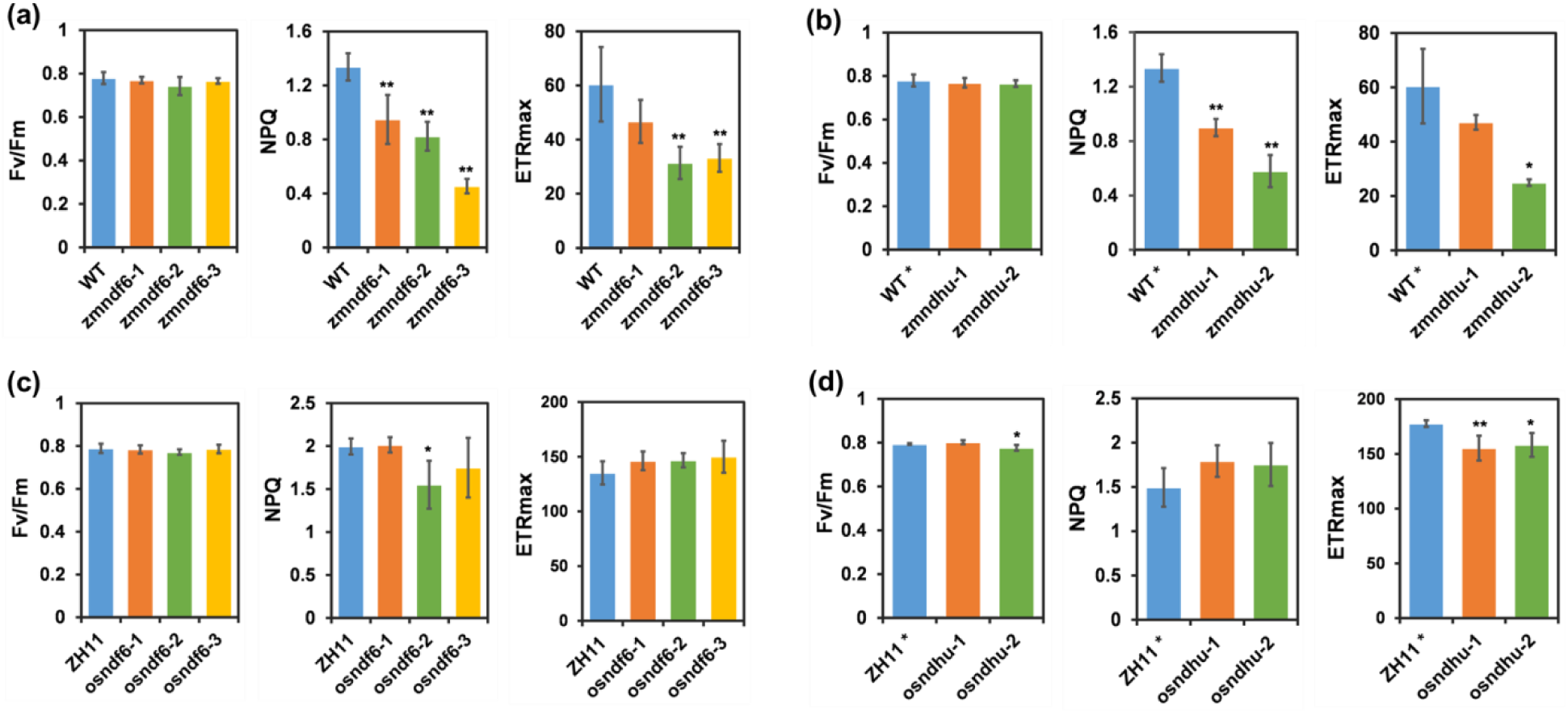
Photosynthetic light reaction parameters in WT and *ndh* mutants. Significant decrease of photosynthetic light reaction parameters in maize but not in rice *ndh* mutants Fv/Fm (maximal PSII quantum efficiency), NPQ (non-photochemical quenching), and ETRmax (maximal electron transport rate) estimated from light response curve. Data are mean ± SE (n = 4 biological replicates). \**P* < 0.05, \*\**P* < 0.01 compared with WT according to Student’s *t* test.

**Fig. S6.**
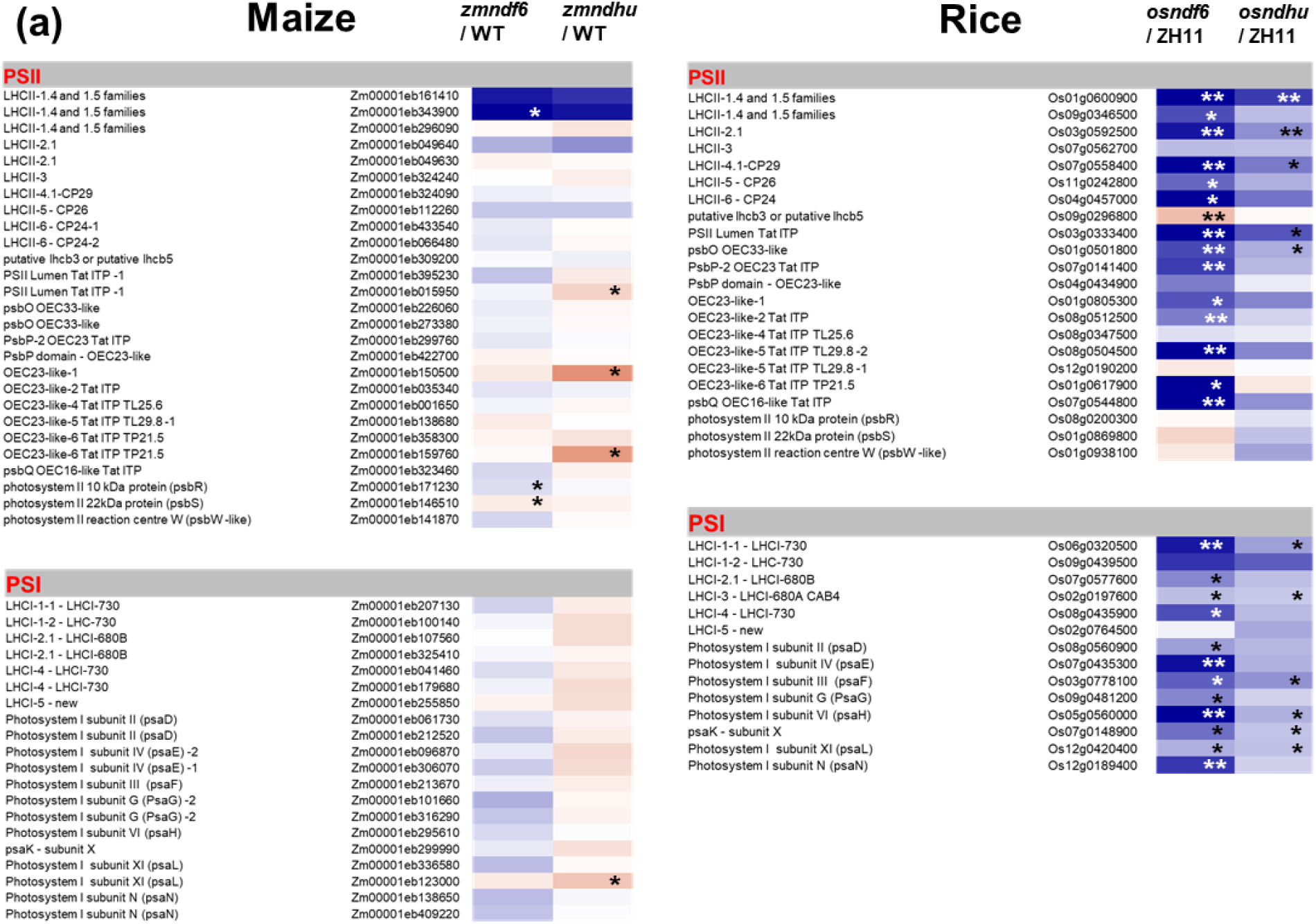
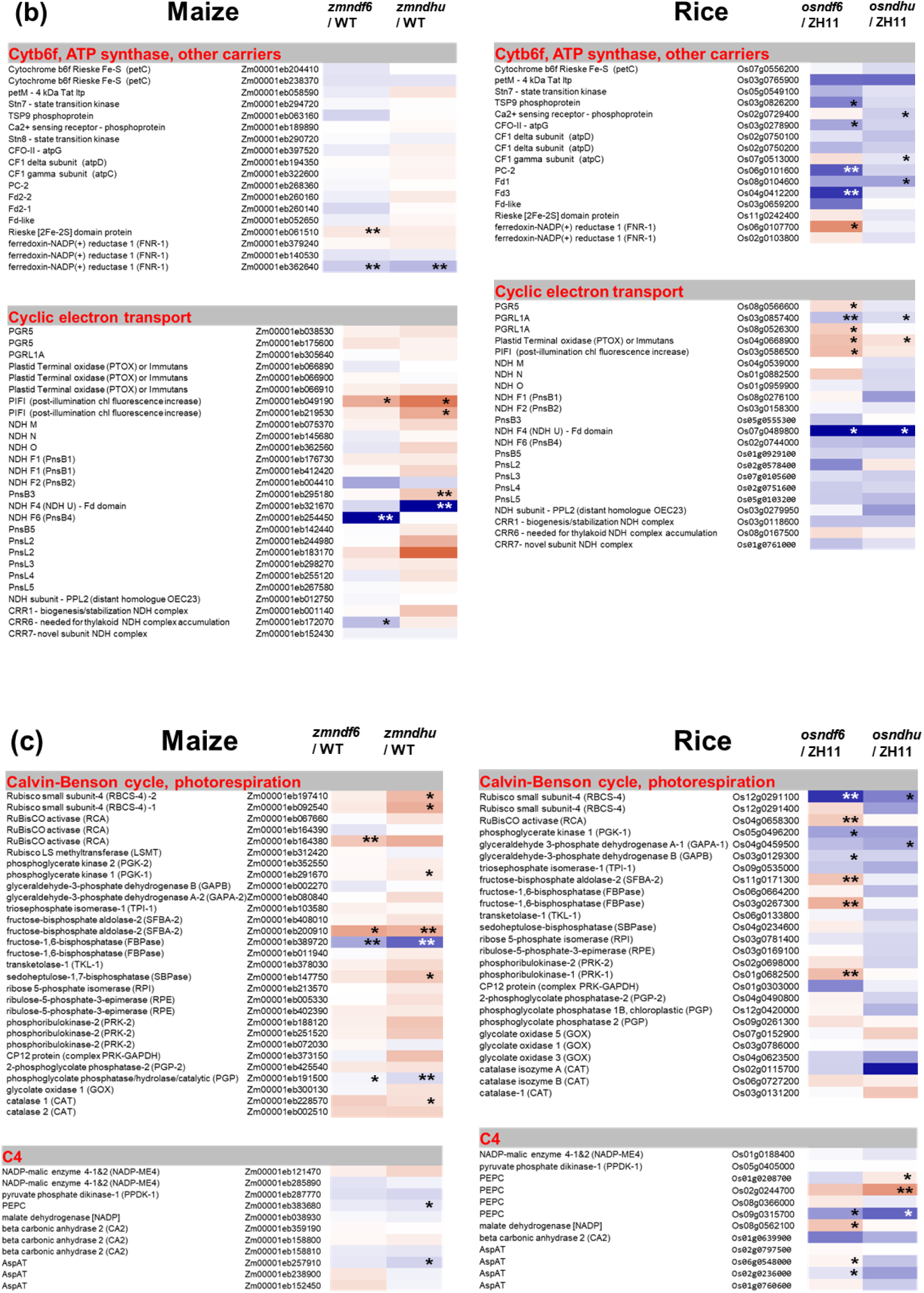
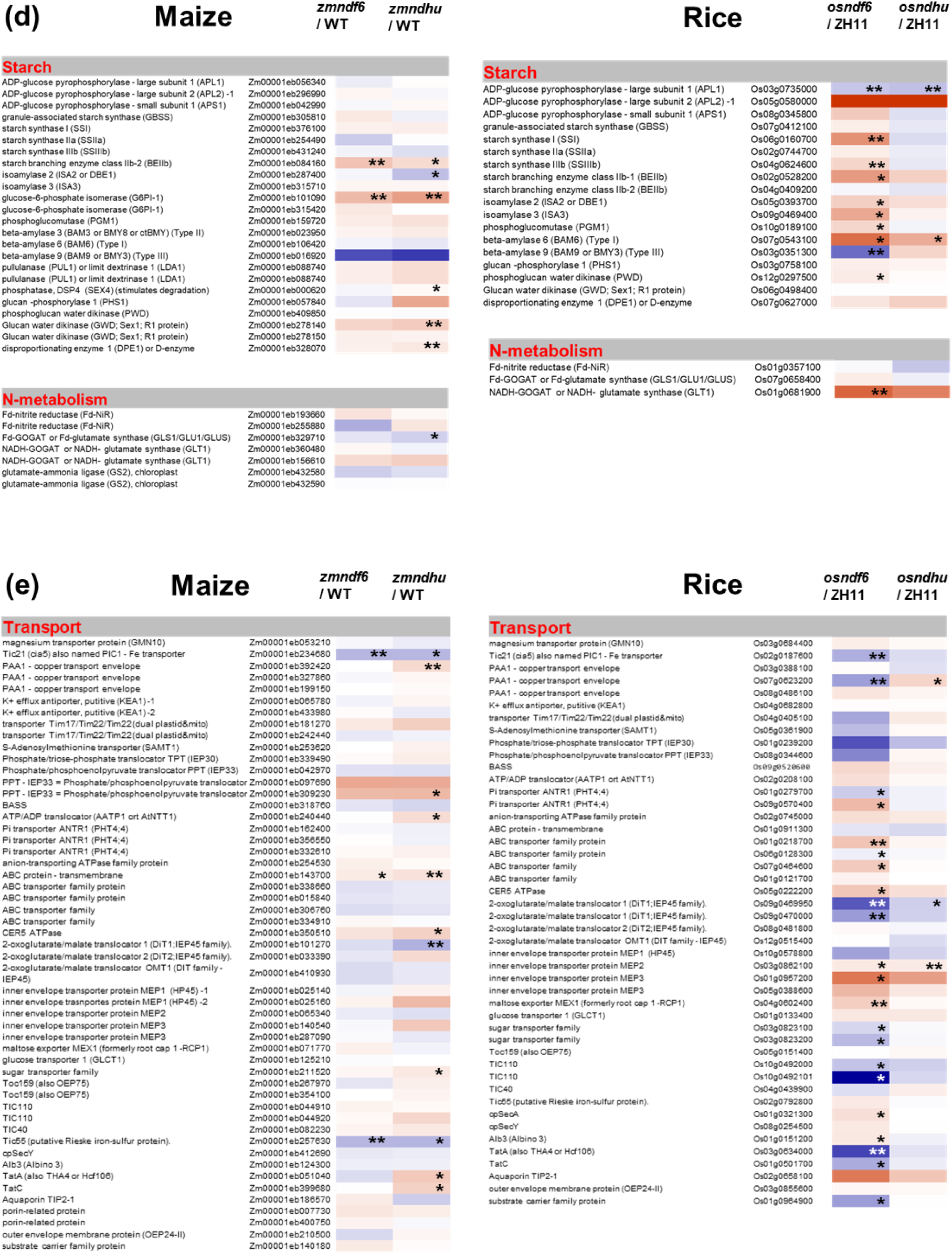
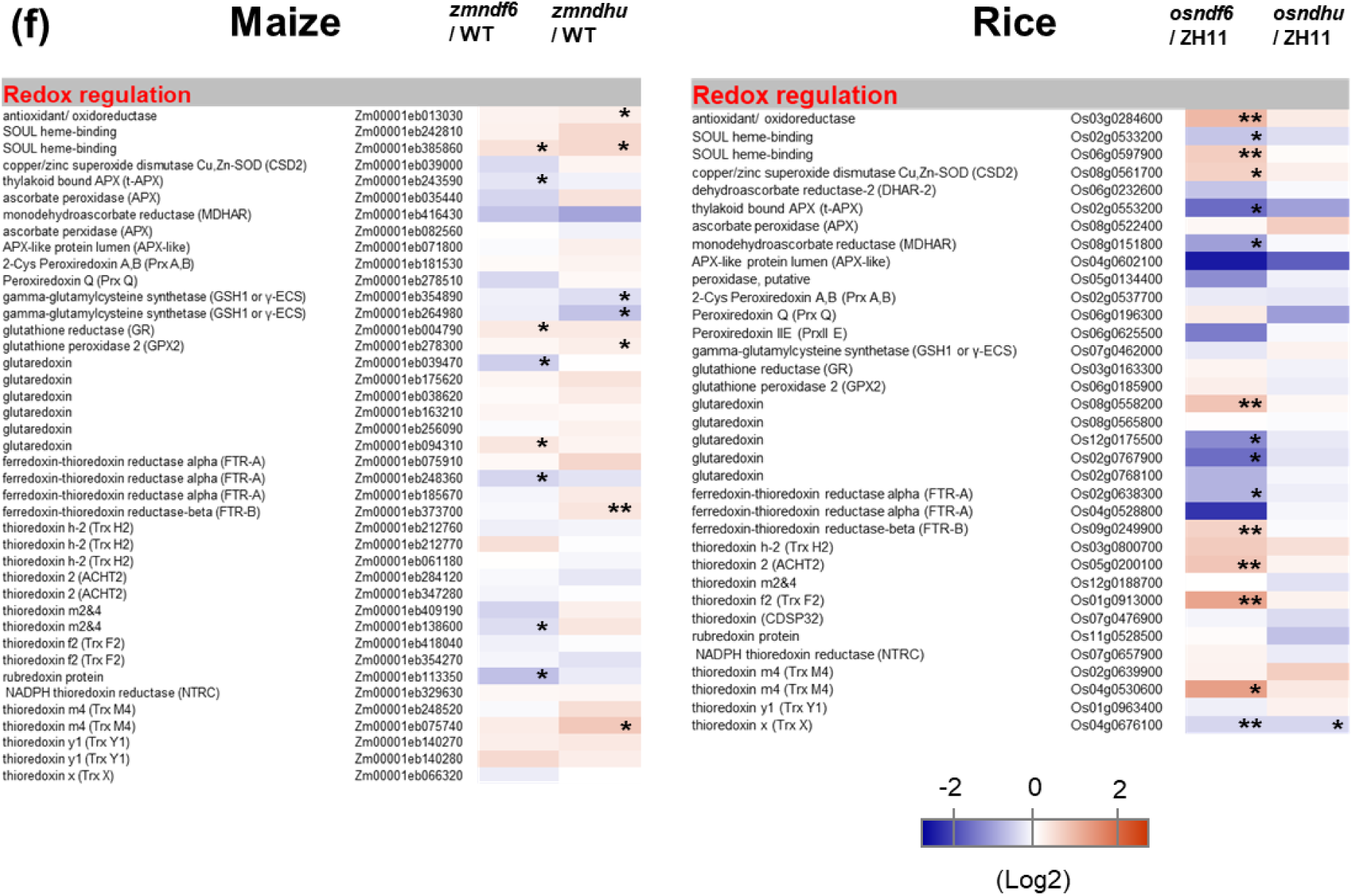
Transcriptome analysis of WT and *ndh* mutant leaves. Transcriptome Analysis of Maize and Rice *ndh* Mutants (a, b) Heatmap shows differentially expressed genes related to photosynthetic linear electron transport (such as PSII, PSI, Cytb6f, ATP synthase) and cyclic electron transport in maize and rice. (c) Expression patterns of genes related to Calvin-Benson cycle, photorespiration, and C_4_ pathway in maize and rice. Other pathways include starch synthesis, N-metabolism (d), transport system (e), and redox regulation (f). Heatmap data represents mean value of 3 biological replicates. *P < 0.05, **P < 0.01 compared with WT according to Student’s t test.

**Fig. S7.**
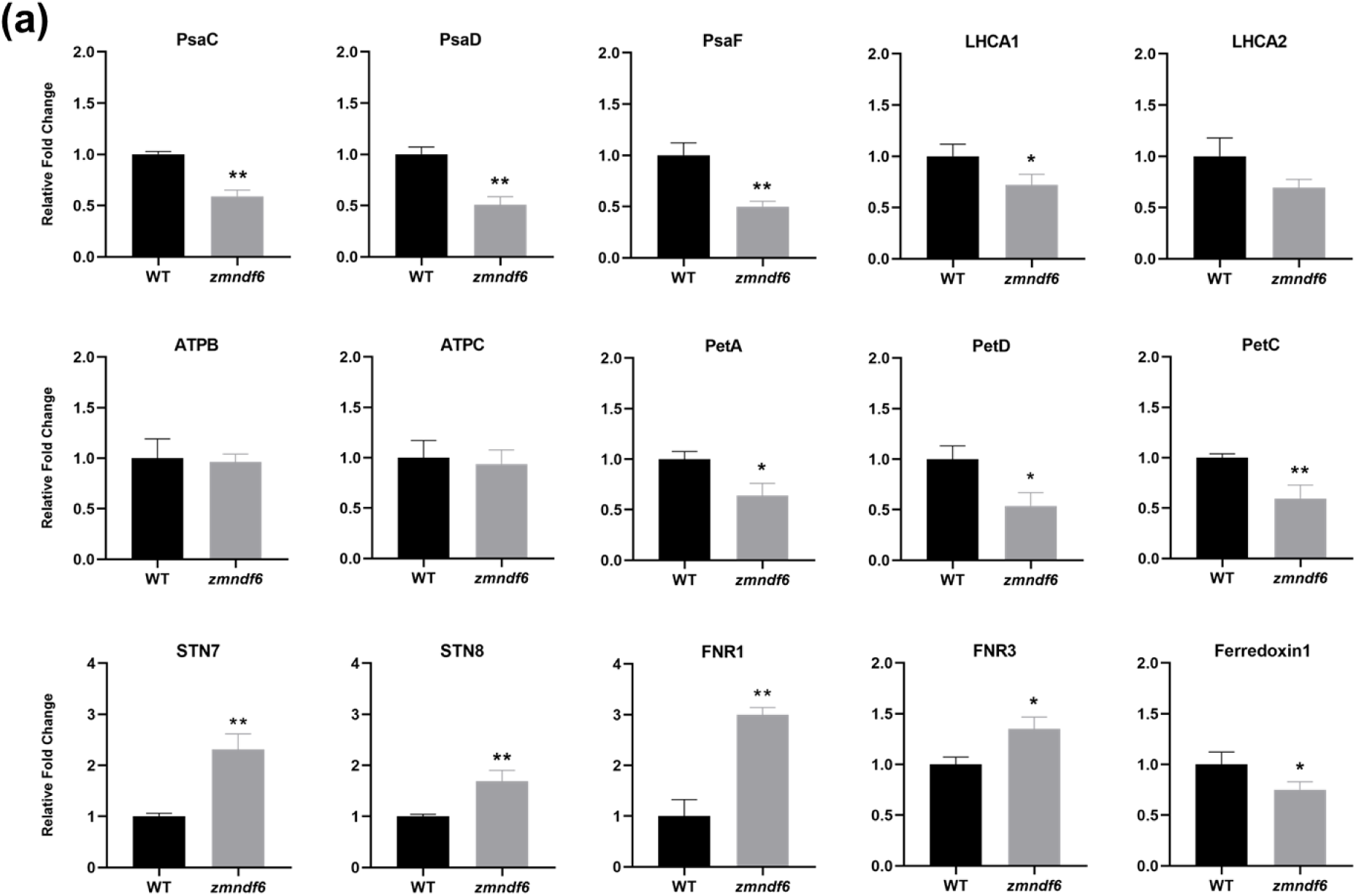
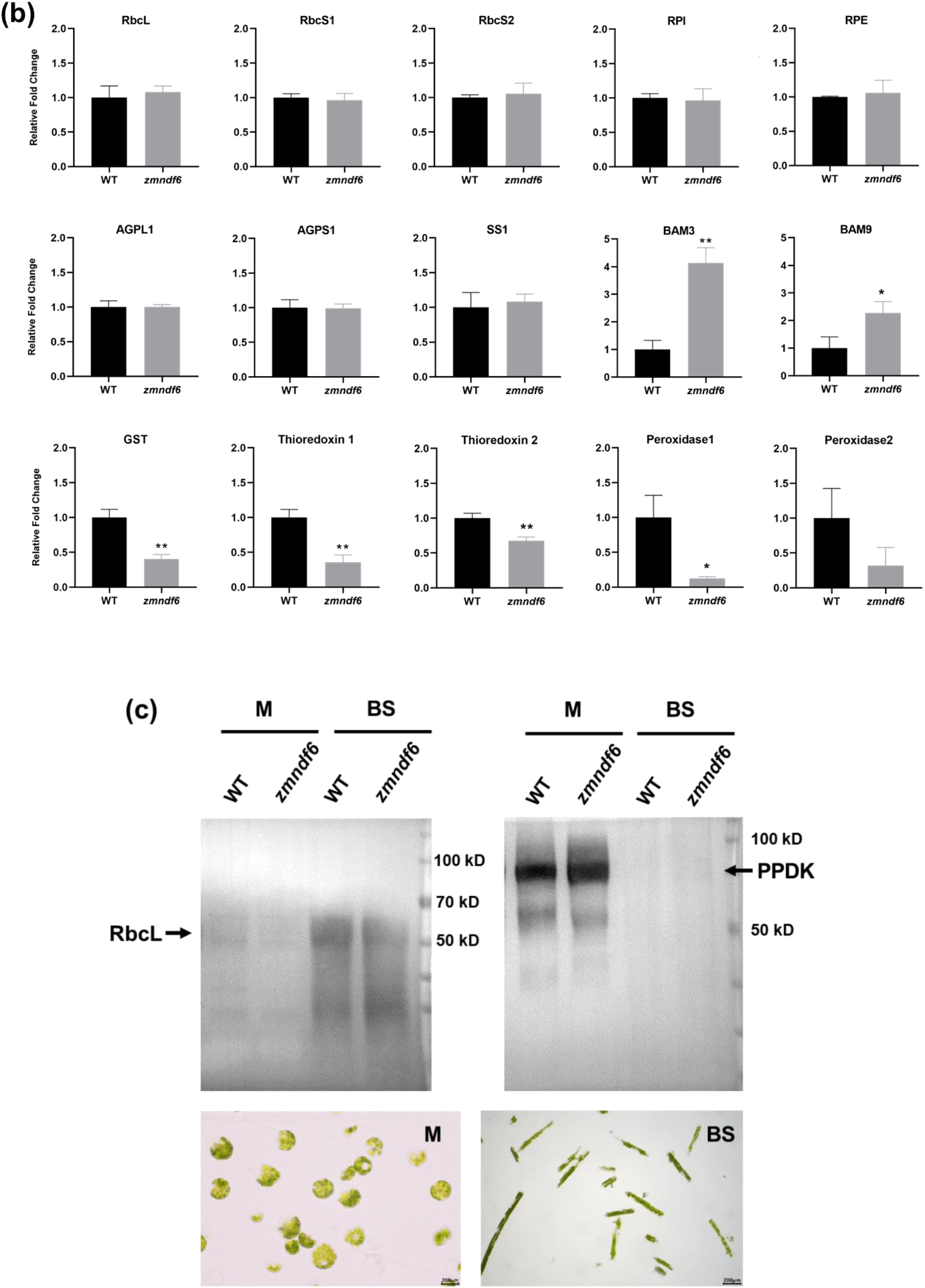
Changes of BS cell protein content in maize *zmndf6* mutants Changes of BS cell protein content related to electron transport (a), metabolism and redox (b) supplementary to Fig 6d. Comparisons were made between WT and *zmndf6* mutant according to quantitative values from BS cell proteomics. Data are mean ± SE (n = 3 biological replicates). \**P* < 0.05, \*\**P* < 0.01 compared with WT according to Student’s t test. (c) Light microscope and immunoblot of RbcL (BS cell enriched) and PPDK (M cell enriched) to test the purity of isolated BS cell samples of WT and *zmndf6* mutant.

**Table S1.** Comparison of the BS/M ratio (mRNA or protein) of selected cyclic electron transport related components in maize (Derived from a, Li et al., 2010; b, Majeran et al., 2008; c, Friso et al., 2010)

| Name | Accession number | Bundle sheath:<br>mesophyll ratio (mRNA) | Bundle sheath:<br>mesophyll ratio (Protein) |  |
| --- | --- | --- | --- | --- |
| NDH subnit F6 (NDF6) | GRMZM2G162233 | 2.21<sup>a</sup> | 7.58<sup>b</sup> | 2.00<sup>c</sup> |
| NDH subnit F4 (NDHU) | GRMZM2G032253 | 4.26<sup>a</sup> | 3.03<sup>b</sup> | 2.94<sup>c</sup> |
| NDH subnit M (NDHM) | GRMZM2G109244 | 2.73<sup>a</sup> | 4.23<sup>b</sup> | 2.10<sup>c</sup> |
| NDH subnit N (NDHN) | GRMZM2G110277 | 3.17<sup>a</sup> | 2.37<sup>b</sup> | 2.13<sup>c</sup> |
| NDH subnit O (NDHO) | GRMZM2G133844 | 3.78<sup>a</sup> | 2.88<sup>b</sup> | 2.74<sup>c</sup> |
| PGR5 | GRMZM2G017045 | -- | -- | 0.90<sup>c</sup> |
| PRGL1A | AC203966.5_FGP002 | -- |  | 1.32<sup>c</sup> |

**Dataset S1** Transcriptome data of maize *zmndf6* and *zmndhu* mutants

**Dataset S2** Transcriptome data of rice *osndf6* and *osndhu* mutants

**Dataset S3** Expression of photosynthetic genes in maize and rice *ndh* mutants

**Dataset S4** BS cell proteomics of maize WT and *zmndf6* mutant

**Dataset S5** Metabolite analysis of maize and rice *ndh* mutants

**Dataset S6** Summary of statistics

**Dataset S7** Summary of primers

